# Histone methyltransferase DOT1L differentially affects the development of dendritic cell subsets

**DOI:** 10.1101/2025.09.05.674429

**Authors:** Rianne G. Bouma, Willem-Jan de Leeuw, Aru Z. Wang, Muddassir Malik, Joeke G.C. Stolwijk, Veronique A.L. Konijn, Anne Mensink, Natalie Proost, Maarten K. Nijen Twilhaar, Tibor van Welsem, Negisa Seyed Toutounchi, Alsya J. Affandi, Jip T. van Dinter, Fred van Leeuwen, Joke M.M. den Haan

## Abstract

Dendritic cells (DCs) are important orchestrators of immune responses. Their development in the bone marrow is controlled by transcription factors, but epigenetic mechanisms remain poorly understood. DOT1L is emerging as a key epigenetic regulator in immune cells. By mapping DOT1L-mediated histone H3K79 methylation in canonical DC subsets, we observed that DOT1L modified common as well as DC subset-specific genes. *In vitro-* or *in vivo-*induced deletion of *Dot1l* followed by *in vitro* cell culture resulted in a decrease in myeloid progenitors and plasmacytoid DCs (pDCs) and an increase in cDC2s, while cDC1s remained unchanged. *In vitro* generated *Dot1l-*KO DCs were unable to produce IFNα upon stimulation. Moreover, transcriptomes of *Dot1l*-KO DC subsets exhibited enrichment of antigen presentation pathways and MHC class II surface levels were upregulated in pDCs. Mechanistically, inhibition of DOT1L linked the observed effects to its methyltransferase activity. Together, our data indicate that in DCs DOT1L differentially affects the development of canonical subsets and suppresses antigen presentation pathways.

## Introduction

Through their ability to integrate the innate and adaptive immune system, dendritic cells (DCs) serve as key regulators of the adaptive immune response (*1,2*). DCs comprise several mature subtypes, and their differentiation trajectory is complex, involving both lymphoid and myeloid lineages. Consequently, DC differentiation is still a field of active study. DCs are commonly divided in conventional DCs (cDCs), including cDC1s and cDC2s, and plasmacytoid DCs (pDCs) (*1–3*). cDC1s have been identified as important stimulators of CD8^+^ T cells through cross-presentation of cell-associated antigens as well as producing IL-12 for T helper 1 cell (Th1) activation. cDC2s have a more prominent role in the activation of Th2, Th17 and B cell responses within the secondary lymphoid organs (*1,4*). The cDC2 subset is further subdivided into the Notch2-dependent T-bet^+^ cDC2A subset that can secrete IL-23 and contribute to germinal center formation and antibody production (*5–8*). In addition, the T-bet^-^ cDC2B subset is transcription factor KLF4-dependent and instigates a Th2 response to parasites and allergens (*6,9*). pDCs are mostly involved in the response to viral infections and contribute to systemic autoimmunity (*3,10,11*) They express toll like receptor (TLR)7 and TLR9 and respond to single stranded RNA or CpG with rapid production of type I interferons (IFNs) (*10*). Through the secretion of type I IFNs, pDCs contribute to the maturation of cDC1s and expansion of CD8^+^ T cells and the regulation of NK cell expansion (*10,12*).

The ontogeny of cDCs and pDCs is subject of ongoing investigation. Both are dependent on the growth factor FMS-like tyrosine kinase 3 ligand (FLT3L) during development and for maintenance. Moreover, their developmental routes have some overlapping as well as distinct features (*1–3*). cDCs develop from the myeloid lineage via Common Myeloid Progenitors (CMPs), Monocyte Dendritic cell Progenitors (MDPs) and Common Dendritic cell Precursors (CDPs). CDPs have a high proliferation rate and consist of multiple pre-disposed subpopulations (*13–15*). Pre-cDC1s, pre-cDC2As and pre-cDC2Bs, developing from CDPs in the bone marrow (BM), migrate to the peripheral tissues and further mature into cDC1s, cDC2As and cDC2Bs (*1,16*). Contributing to the cDC2 lineage are pDC-like cells, also known as transitional DCs, expressing pDC-associated markers which have recently been identified as pre-cDC2A cells (*17–20*). Two main developmental routes have been described for pDCs. The shared dependence on FLT3L and the identification of shared precursors initially indicated that pDCs developed from the myeloid lineage similar to cDCs (*21–23*). However, a lymphoid lineage origin for pDCs in which they develop from Common Lymphoid Precursors (CLPs) has recently been established (*24–27*). Regardless of their ontogeny, pDCs complete their full maturation process in the BM, after which they migrate to the tissues. Finally, the DC3 subset is the most recent identified DC subtype and constitutes an independent lineage that derives from Ly6C^+^ MDPs (*28*).

While transcriptional regulation of DC differentiation has been studied extensively, the role of the epigenome in DC differentiation is still poorly understood (*29–31*). At the level of genome folding, recently, a role in cDC1 development and function was found for the cohesion complex (*32*). At the level of the nucleosome, deficiency of the histone lysine demethylase KDM5C stimulates the development of cDC1s and cDC2Bs, and leads to increased numbers of non-IFN-producing pDCs with increased inflammatory gene expression (*33*). Additionally, deficiency of the histone deacetylases HDAC1 and HDAC3 impairs pDC and cDC2 development, while cDC1 development is not affected (*34–36*). However, HDAC1 and HDAC3 are enzymes with broad substrate specificities (*37*) and may therefore affect DCs by mechanisms beyond erasing acetyl marks from chromatin. In thymocytes, HDAC1 has been shown to crosstalk with the chromatin modifier Disruptor of telomeric silencing 1 – like (DOT1L) (*38*). This histone methyltransferase is emerging as an important gatekeeper in the differentiation and functioning of macrophages, NK, B and T cells (*38–47*). Recent studies suggest that DOT1L affects DC maturation and tolerance induction, although its role in DC subset differentiation remains unknown (*31,48,49*).

DOT1L has one canonical substrate and is the sole methyltransferase of histone 3 lysine 79 (H3K79) (*50–52*). H3K79me is predominantly deposited in the promotor-proximal gene-body regions of transcribed genes (*53*). DOT1L facilitates the mono-(me1) di-(me2) and tri-methylation (me3) of H3K79, and its activity is enhanced through transcriptional elongation and transcription-associated histone modifications (*51,54*). In agreement with this co-transcriptional regulation, the level of H3K79 methylation (H3K79me) correlates with the level of gene expression (*51,52,55,56*). Highly specific inhibitors have been developed that specifically block DOT1L catalytic activity, lead to loss of H3K79me, and have potential therapeutic applications (*57–62*). However, H3K79 methylation is remarkably stable and mainly lost through cell division (*61,62*), which is agreement with the absence of a demethylase known to act on this mark (*41,43*). While DOT1L activity mainly seems restricted towards H3K79me, cellular functions of DOT1L independent of its methyltransferase activity have been described as well (*63–68*).

Here, we characterized the role of DOT1L in the differentiation of DCs both *in vitro* and *in vivo*. We show that genetic ablation of *Dot1l* followed by *in vitro* cell culture of BM cells leads to a decrease in myeloid precursors and pDCs, and an increase in cDC2s. This role of DOT1L is linked to its catalytic activity, as chemical inhibition recapitulated these findings. In classical DC subsets, we confirmed that dimethylation of H3K79 (H3K79me2) is deposited in a canonical pattern across gene bodies of transcribed genes. The targets of DOT1L include genes encoding transcription factors involved in pDC differentiation (*Tcf4, Bcl11a, Irf8 and SpiB*) and the expression of these transcription factors was decreased in *Dot1l*-KO pDCs *in vivo*. In addition, antigen presentation-related pathways were enriched in the transcriptomes of all *Dot1l-*KO DC subsets. Combined, these results suggest that DOT1L plays an essential role in the function and differentiation of DCs, especially in pDCs. Our quantitative H3K79me2 epigenomics map, combined with perturbation and transcriptomic data provide an important resource for gaining more insights into how DC development and function is guided by epigenetic modulation.

## Methods

### Mice

Cre-ER^T2^ C57BL/6 mice contained Cre fused to the human estrogen receptor (Cre-ER^T2^) integrated at the ROSA-26 locus, as described elsewhere (*69*). This strain was crossed with Dot1l^fl/fl^ mice from the Dot1ltm1a(KOMP)Wtsi line as generated by the Wellcome Trust Sanger Institute (WTSI) and obtained from the KOMP Repository (www.komp.org) (*70*) and as described previously to generate the Cre-ERT2;*Dot1l*^fl/fl^ strain (*38,41*). Cre-ER^T2^;*Dot1l*^fl/fl^;OT-I mice were generated by crossing Cre-ER^T2^;*Dot1l*^fl/fl^ mice with OT-I (B6J carrying the OT-1 T cell receptor transgenes) mice (kindly gifted by the Ton Schumacher group, originally from Jackson labs). Cre-ER^T2^;*Dot1l^wt/wt^*(WT) and Cre-ER^T2^;*Dot1l^fl/fl^* mice were used for experiments, without pre-selection based on OT-I presence. Experiments were performed in-house at the animal facility of the NKI. The mice were between 8 weeks and 8 months old and matched for age and gender. To induce the genetic ablation of DOT1L *in* vivo, Cre-ERT2;*Dot1l^wt/wt^* mice or Cre-ERT2;*Dot1l^fl/fl^* mice were injected intra-peritoneally with 75mg/kg tamoxifen (Sigma Aldrich, an overview of key reagents used in this study can be found in Supplementary Table S1) for three consecutive days. Experiments were performed according to institutional, national and European guidelines as approved by the Animal Ethics Committee of the NKI (CCD number: AVD30100202215981).

### BM cultures

Murine BM cells (from C57BL/6 mice, Cre-ER^T2^;*Dot1l^wt/wt^*mice or Cre-ER^T2^;*Dot1l^fl/fl^* mice; indicated in-text) were isolated from femurs and tibias by flushing the bones using a 10 mL syringe and 25G needle. The cells were filtered through a 70 μm cell strainer and concentrated by centrifugation at 1200 rpm for 10 minutes with low break. Next, the cells were cultured at a concentration of 1 million cells per mL in 2 mL DC medium. DC medium consisted of RPMI-1640 (Gibco) supplemented with 10% heat-inactivated FCS (Biowest), 50 U/mL penicillin (Lonza, Basel, Switzerland), 50 µg/mL streptomycin (Lonza), 50 µM β-mercaptoethanol (Gibco) and 100x diluted Glutamax (Gibco), as well as 200 ng/mL FLT3-L (Peprotech) and 50 ng/mL SCF (Peprotech). On day 3 after the start of the cultures, each well was divided over 2 wells and 1 mL fresh DC medium without SCF was added. On day 7, the cells were harvested, and each well was washed with 1 mL PBS 2 times. After centrifugation for 5 minutes at 1500 rpm, the cells were stained using a flow cytometry panel and acquired on the Aurora spectral flow cytometer (Cytek, Amsterdam, the Netherlands). Where indicated, tamoxifen or DOT1L inhibitor was added to the cultures. 4-hydroxy-tamoxifen (Sigma Aldrich) was added on day 0 at a concentration of 10 nM. DOT1L inhibitor SGC-0946 (Selleckchem) was added on day 0, 3 and 5 at a concentration of 1 μM. The same volume of DMSO was added to the control wells.

### H3K79me2 staining and Dot1l PCR

BM suspensions were prepared as described above. Splenocytes were isolated by meshing spleens three times through a 70 µm strainer (BD Falcon, Franklin Lakes, NJ, USA) to create single cell suspensions. The samples were incubated with lysis buffer (150 mM NH_4_Cl, 10 mM KHCO_3_, 0.2 mM EDTA) for three minutes on ice to lyse erythrocytes. For flow cytometry, 2.5×10^5^ cells per sample were washed with PBS and subsequently stained with Zombie-NIR (BioLegend). Intracellular staining was performed using the Transcription Factor Buffer kit (BD Biosciences), according to manufacturer’s protocol. Cells were stained with anti-H3K79me2 (Millipore) in 0.25 % Sodium Dodecyl Sulfate (SDS) supplemented perm/wash buffer (BD Biosciences) followed by 1:1000 secondary antibody staining (Goat-anti-Rabbit AF488; Invitrogen) as described previously (*41*). Samples were measured using the LSR-Fortessa II (BD Biosciences) and analyzed using FlowJo V10.9.0 (BD Biosciences).

To confirm the deletion of exon 2 of *Dot1l* by PCR, genomic DNA was isolated using the ISOLATE II Genomic DNA Kit (Bioline) as per manufacturer’s protocol. *Dot1l^WT^, Dot1l^fl^,* and *Dot1l^Δ^* (exon 2) alleles were detected as described previously (*41*), using the following primers; Dot1l_FWD: GCAAGCCTACAGCCTTCATC, Dot1l_REV: CACCGGATAGTCTCAATAATCTCA and Dot1l_Δ: GAACCACAGGATGCTTCAG. PCR reactions were performed using MyTaq Red Mix (GC Biotech). Agarose gel electrophoresis was performed to determine the genotype. The efficacy of floxing out exon 2 was determined by quantifying band densities using ImageJ (version 1.54g), correcting for product size and dividing the band density of the *Dot1l^Δ^* band over the total *Dot1l* band density within the sample.

### Sample preparation of in vivo experiments

Bone marrow and spleen suspensions were prepared from Cre-ER^T2^;*Dot1l^wt/wt^* mice or Cre-ER^T2^;*Dot1l^fl/fl^*mice. Spleens were mechanistically dissociated and digested with 4 mg/mL lidocaine (Sigma Aldrich), 4 Wunsch units/ml Liberase TL (Roche) and 50 mg/mL DNase I (Roche) for 20 minutes at 37 °C with continuous stirring. Digestion was then halted by addition of cold medium (RPMI-1640 (Gibco) with 10% heat-inactivated FCS (Biowest), 10 mM ethylenediaminetetra-acetic acid (EDTA), 20 mM HEPES and 50 µM 2-mercaptoethanol) and another 10 minutes stirring at 4 °C. Ammonium-chloride-potassium lysis buffer was added to lyse the red blood cells and cells were filtered through a 70 to 100 μm cell strainer immediately afterwards. The cells were then counted and a flow cytometry staining as well as functional experiments were performed.

### Flow cytometry panels

To identify mature DC subsets in the BM as described in Figure 1, we first incubated the cells with 10 µg/mL of anti-CD16/32 (clone 2.4G2, in-house production) and Fixable Viability Dye eFluor 780 (eBioscience, San Diego, CA, USA). Next, we stained the cells for 20 minutes at 4 °C with the following combination of antibodies: anti-XCR1 (Clone ZET; Biolegend), anti-MHC class II (Clone M5/114.15.2; Biolegend), anti-CD11c (Clone N418; Biolegend), anti-BST2 (Clone 927; Biolegend), anti-CD3 (Clone 17A2; Biolegend), anti-CD19 (Clone 6D5; Biolegend), anti-NK1.1 (Clone PK136; Biolegend), anti-Siglec-H (Clone REA819; Miltenyi Biotec) and anti-Sirpα (Clone P84; Biolegend). Finally, the cells were fixed using 2 % PFA (Electron Microscopy Sciences, Hatfield, PA, USA) and subsequently acquired on the Aurora spectral flow cytometer (Cytek).

**Figure 1.**
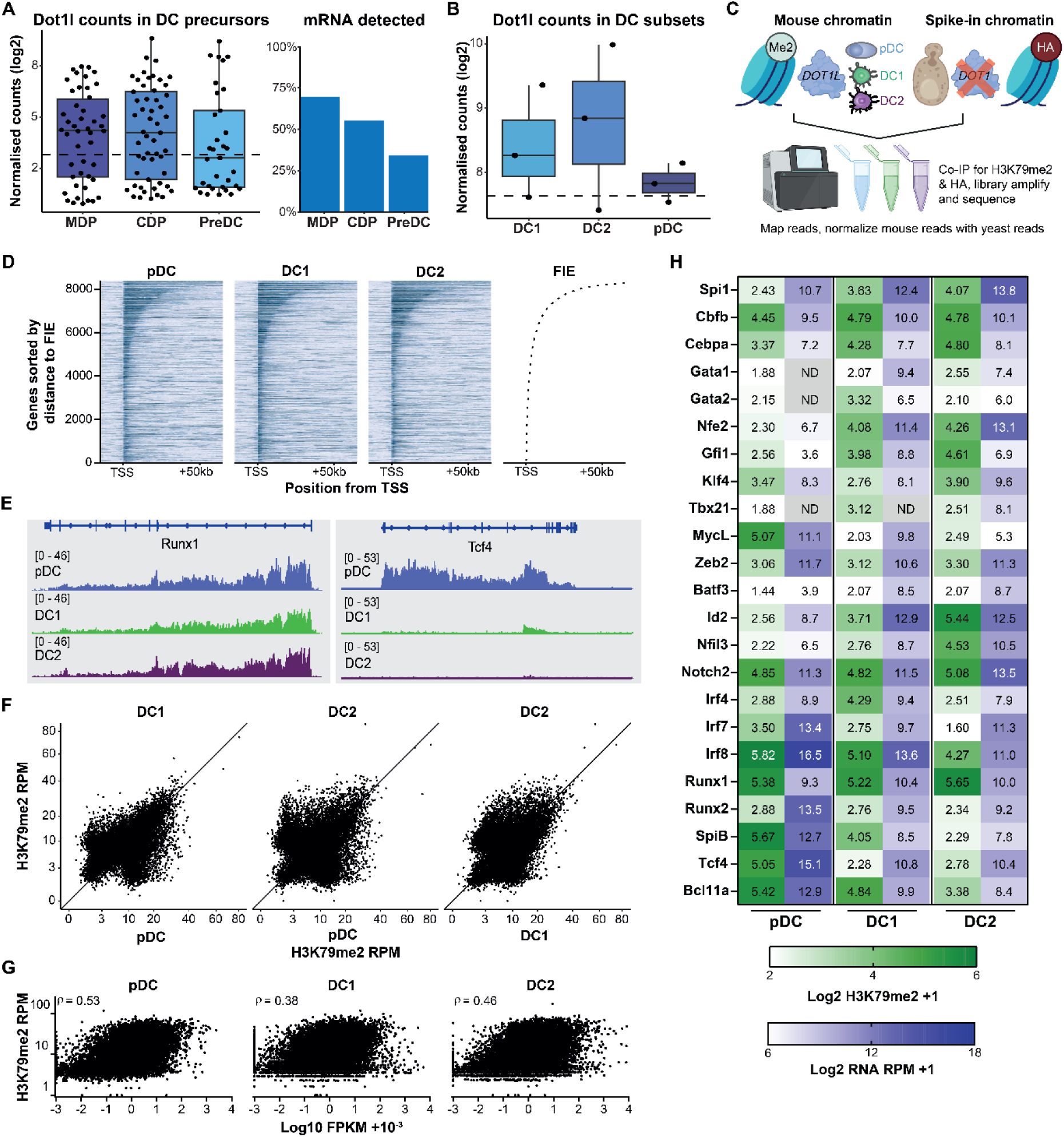
DOT1L H3K79me2 methylation in DC subsets. **(A)** *Dot1l* counts derived from scRNA sequencing analysis of sorted DC precursors. Left: median of all genes is plotted as dashed line. Right: percentage of cells with *Dot1l* counts higher than 0 (*Dot1l* detected) versus cells with no *Dot1l* counts (not detected). Dataset acquired from Schlitzer *et al.* (2015 (*13*). **(B)** *Dot1l* counts derived from bulk RNA sequencing analysis of sorted DC subsets. The median of all genes is plotted as dashed line. Dataset acquired from Rodrigues *et al.* (2023) (*17*). **(C)** Spike-in normalized H3K79me2 ChIPseq workflow. Sorted Siglec-H^pos^ BST2^pos^ pDC, CD11c^pos^ XCR1^pos^ cDC1 and CD11c^pos^ sirpα^pos^ cDC2 pools (N=2 biological replicates) were spiked with chromatin from a *dot1Δ* and HHT2-HA tagged yeast strain and co-immunoprecipitated for H3K79me2 and HA in parallel. Reads mapped to the *S. cerevisiae* genome were used to normalize the H3K79me2 reads within each sample. **(D)** Tornadoplot showing H3K79me2 signal in relation to the transcription start site (TSS) and first internal exon (FIE). For each subset, the top 50% active genes based on expression are displayed, ranked according to distance to the FIE (short to long), as displayed in the rightmost graph. **(E)** Snapshot of the normalized H3K79me2 data obtained from **(C)** for two representative genes. Data was visualized using *IGV*. **(F)** Dot plots comparing H3K79me2 peaks between the sorted DC subsets. **(G)** Dot plot correlating H3K79me2 signal near the TSS (N=2) and RNA expression (N=1) for each sorted DC subset. Expression data was normalized for transcript length by calculating the Fragments Per Kilobase of transcript per Million mapped reads (FPKM) values for each gene. The calculated Spearman’s rank correlation coefficients are displayed within each graph. **(H)** Heatmap of selected transcription factors, quantifying H3K79me2 signal near the TSS (green gradient) and expression based on RNAseq (blue gradient) in the sorted subsets ND = not detected.

To identify precursor cell subsets in the BM as described in Figure 2, 3 and 4, single cell suspensions were incubated with 10 µg/mL of anti-CD16/32 (clone 2.4G2, in-house production) and LIVE/DEAD Fixable Blue Dead Cell Stain (Thermo Scientific, Waltham, MA, USA). Afterwards, the cells were stained with the following antibody panel for 20 minutes at 4 °C: anti-MHC class II (Clone M5/114.15.2; Biolegend), anti-CD117 (Clone 2B8; Biolegend), anti-CD11c (Clone N418; Biolegend), anti-BST2 (Clone 927; Biolegend), anti-CD3 (Clone 145-2C11; Immunotools), anti-NK1.1 (Clone PK136; Immunotools), anti-Erythroid cells (Clone TER-119; Immunotools), anti-CD90 (Clone MRC OX-7; Immunotools), anti-ly6G (Clone RB6-8C5; Immunotools), anti-B220 (Clone RA3-6B2; Biolegend), anti-CD135 (Clone A2F10; Invitrogen), anti-CD127 (Clone A7R34; Biolegend), anti-CD19 (Clone 6D5; Biolegend), anti-CD115 (Clone AFS98; Biolegend), anti-Sirpα (Clone P84; Biolegend), anti-Siglec-H (Clone REA819; Miltenyi Biotec) and anti-Ly6D (Clone REA906; Miltenyi Biotec). The cells were subsequently fixed using 2 % PFA (Electron Microscopy Sciences) and acquired on the Aurora spectral flow cytometer (Cytek).

**Figure 2.**
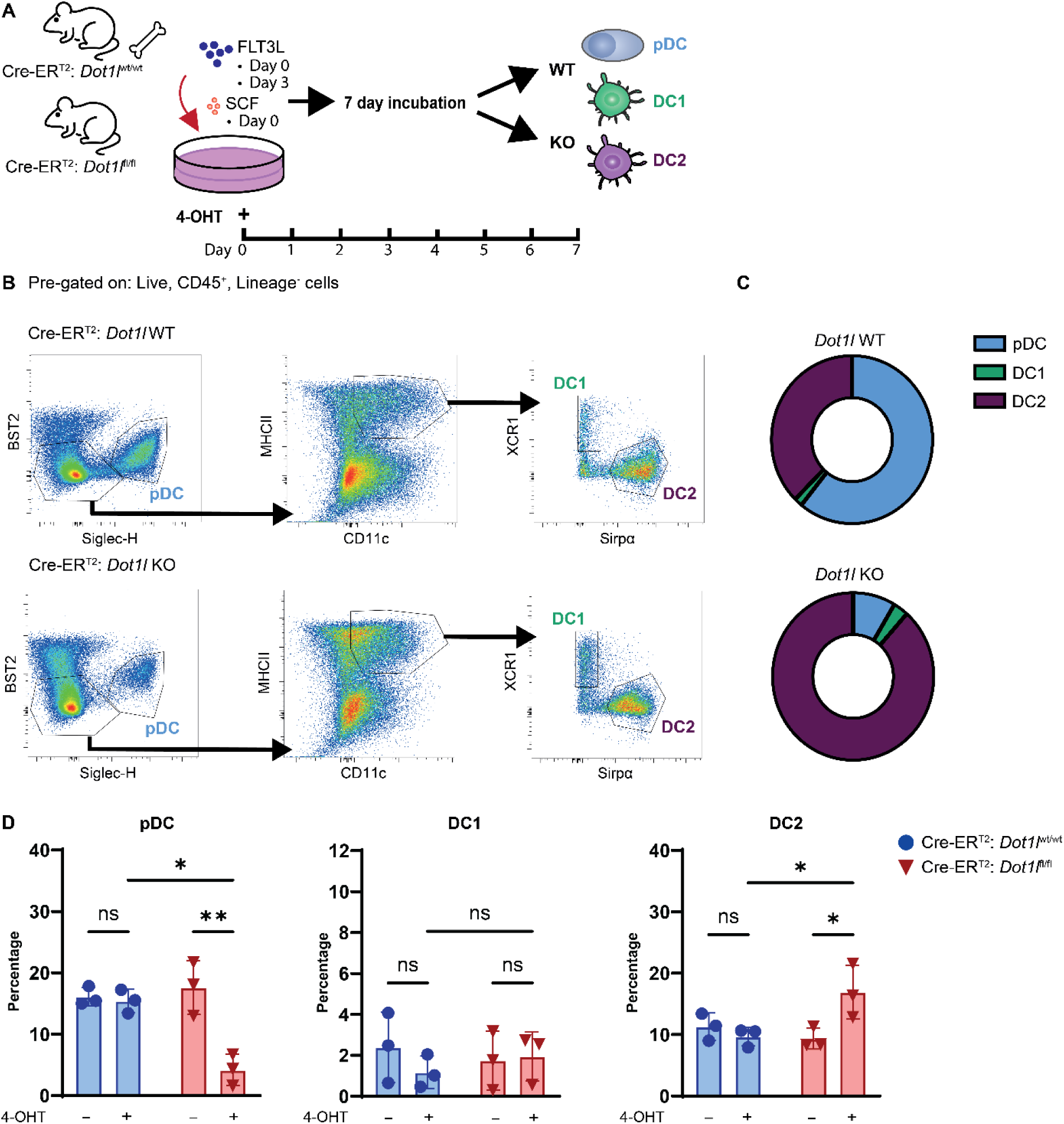
Decreased development of pDCs and increased development of cDC2s upon *in vitro* deletion of *Dot1l* in FLT3L/SCF BM cultures. **(A)** Schematic overview of the setup of the 7-day BM culture. Total BM of Cre-ER^T2^ *Dot1l*^wt/wt^ and Cre-ER^T2^ *Dot1l*^fl/fl^ mice was cultured with or without 4-hydroxytamoxifen and with SCF and FLT3L to facilitate the development of DCs. **(B)** Overview of the gating strategy on total BM SCF-, FLT3L-cultures on day 7. The top row is representative of a *Dot1l* WT sample, and the bottom row is representative of a *Dot1l* KO sample. **(C)** DC subset percentages of Live CD45^pos^ Lineage^neg^ cells. Circle plots represent *Dot1l* WT (Cre-ER^T2^;*Dot1l*^wt/wt^ + 4-hydroxytamoxifen) and *Dot1l* KO (Cre-ER^T2^;*Dot1l*^fl/fl^ + 4-hydroxytamoxifen) conditions. **(D)** Comparison of DC subset percentages of Live CD45^pos^ Lineage^neg^ cells in Cre-ER^T2^ *Dot1l*^wt/wt^ and Cre-ER^T2^ *Dot1l*^fl/fl^ conditions, with and without 4-hydroxytamoxifen to induce KO of *Dot1l*. The individual dots represent the average of technical replicates in one independent experiment, with a total of three independent experiments performed. Error bars indicate mean ± SD. *p < 0.05**p < 0.01, ***p < 0.001, ****p < 0.0001.

**Figure 3.**
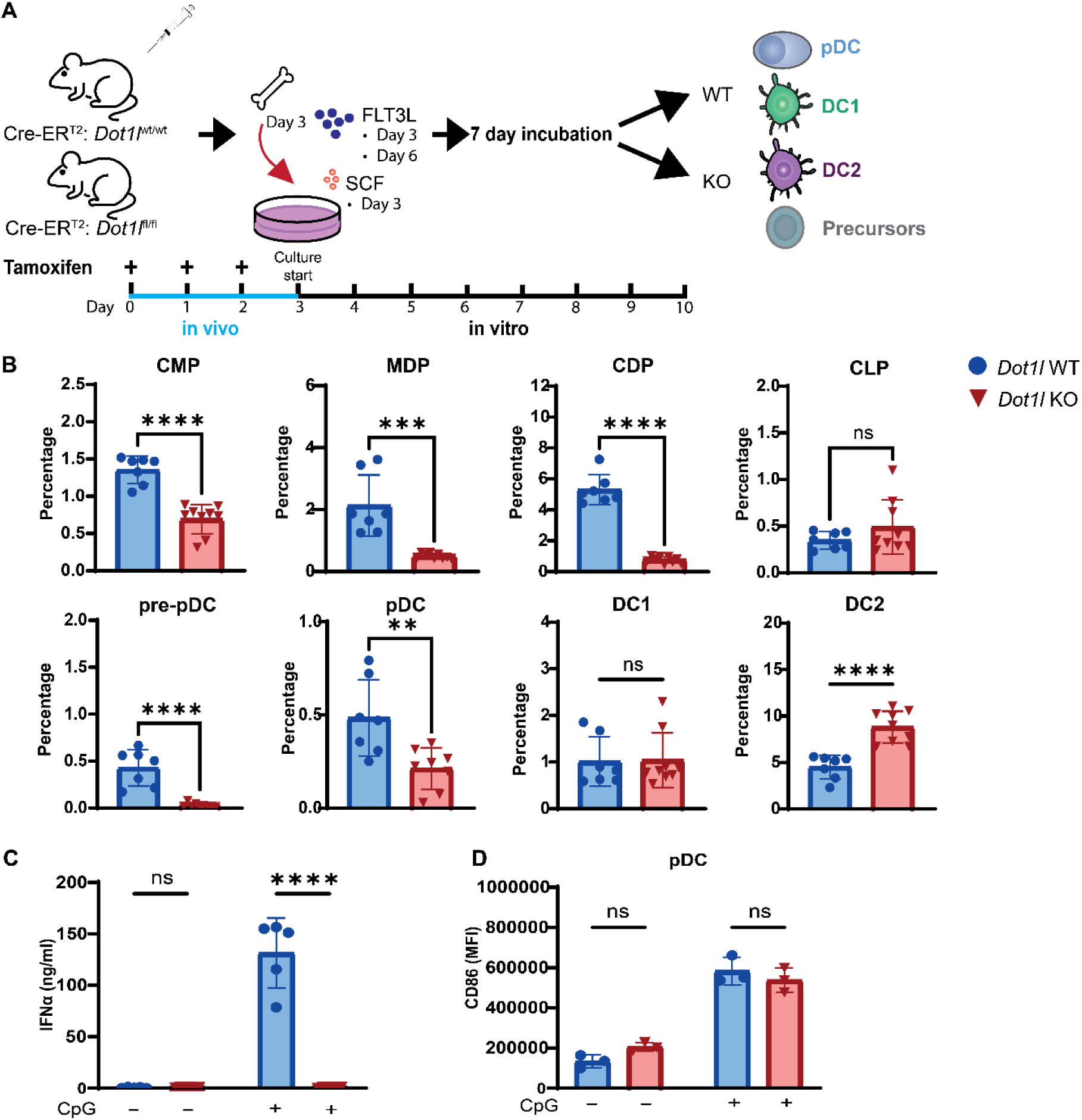
Reduced frequency of pDCs, and myeloid, but not lymphoid precursors after *in vivo* deletion of *Dot1l* and subsequent seven-day SCF-, FLt3L-BM culture (A) Cre-ER^T2^ *Dot1l*^wt/wt^ and Cre-ER^T2^ *Dot1l*^fl/fl^ mice were injected three times with tamoxifen. Subsequently, BM was cultured with SCF and FLT3L. Seven days after the start of the culture, DC and precursor subset composition was evaluated. **(B)** Comparison of myeloid and lymphoid precursor percentages of Live Lineage^neg^ AF^neg^ cells in *Dot1l* WT (Cre-ER^T2^ *Dot1l*^wt/wt^) and *Dot1l* KO (Cre-ER^T2^ *Dot1l*^fl/fl^) conditions. CMPs, MDPs and CDPs were gated from MHC class II^neg^ CD11c^neg^ CD135^pos^ cells and defined as CD117^hi^ CD115^neg^ CMPs, CD117^int^ CD115^hi^ MDPs and CD117^neg^ CD115^hi^ CDPs. CLPs were defined as MHC class II^neg^ CD11c^neg^ CD135^neg^ CD127^pos^. pre-pDCs were gated as CD117^neg^ CD135^pos^ CD11c^pos^ Siglec-H^pos^ Ly-6D^pos^, pDCs as B220^pos^ Siglec-H^pos^ BST2^pos^ and cDC1s and cDC2s as B220^neg^ MHC class II^hi^ CD11c^hi^ Sirpα^neg/pos^. **(C)** On day 7 after the start of the culture, BM cells were stimulated with Class A CpGs (ODN1585) overnight. IFNα levels were determined using ELISA. **(D)** After Class A CpG stimulation, cells were stained for maturation markers. Conditions are shown with and without the addition of CpGs. The individual dots represent individual mice with a total of three independent experiments performed. Error bars indicate mean ± SD. *p < 0.05, **p < 0.01, ***p < 0.001, ****p < 0.0001.

**Figure 4.**
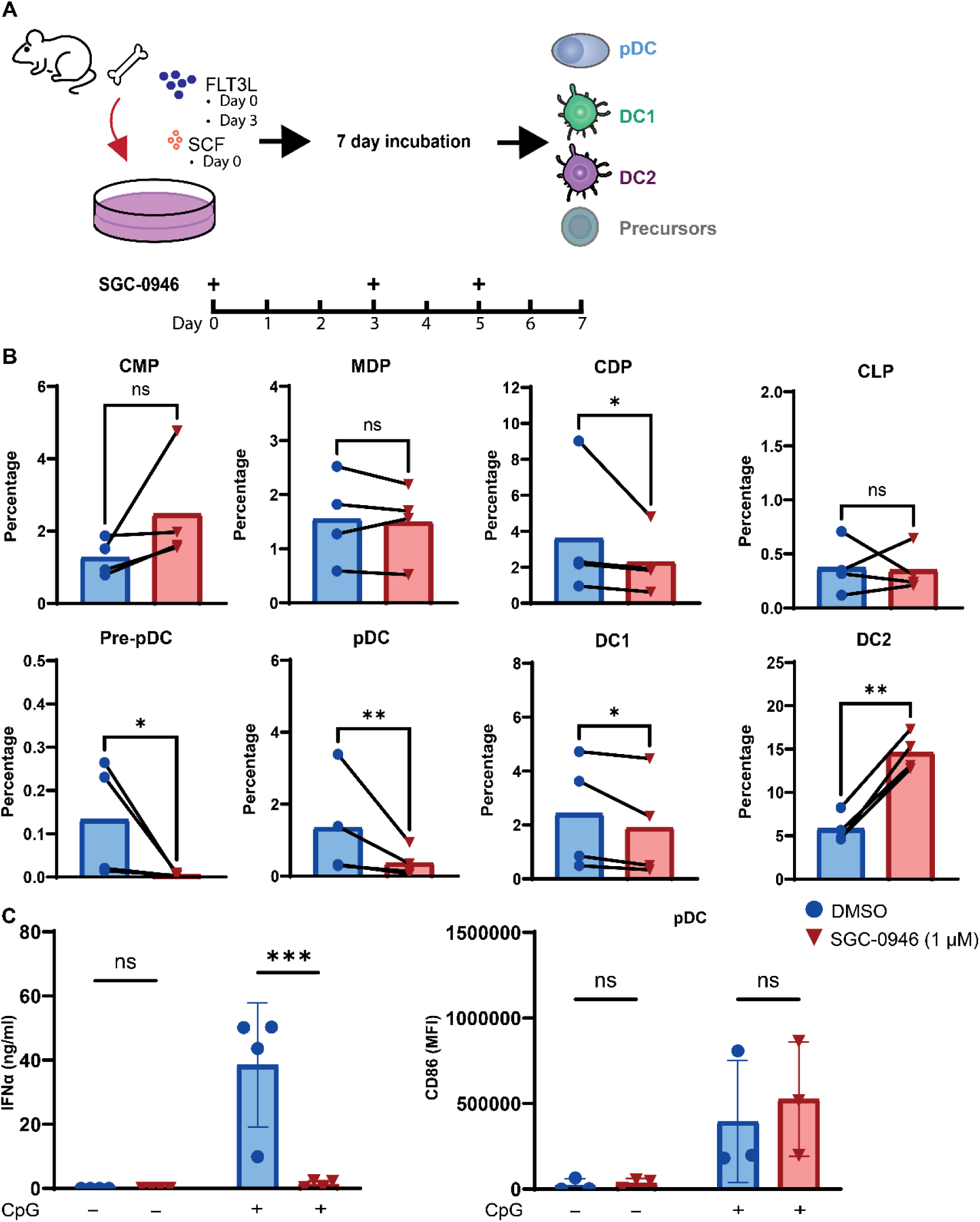
*In vitro* inhibition of DOT1L in WT BM cultures results in decreased generation of pDC and increased generation of cDC2. **(A)** Schematic overview of the setup of the 7-day BM culture. Total BM of WT C57BL/6 mice was cultured with SCF and FLT3L to facilitate the development of DCs. DOT1L inhibitor SGC-0946 (1 µM) was added at day 0, 3 and 5 to ensure constant inhibition of DOT1L during DC development. DMSO (0.1%) was used as a vehicle control. **(B)** Comparison of myeloid and lymphoid precursor percentages of Live Lineage^neg^ AF^neg^ cells in cultures with and without DOT1L inhibitor. CMPs, MDPs and CDPs were gated from MHC class II^neg^ CD11c^neg^ CD135^pos^ cells and defined as CD117^hi^ CD115^neg^ CMPs, CD117^int^ CD115^hi^ MDPs and CD117^neg^ CD115^hi^ CDPs. CLPs were defined as MHC class II^neg^ CD11c^neg^ CD135^neg^ CD127^pos^. pre-pDCs were gated as CD117^neg^ CD135^pos^ CD11c^pos^ Siglec-H^pos^ Ly-6D^pos^, pDCs as B220^pos^ Siglec-H^pos^ BST2^pos^ and cDC1s and cDC2s as B220^neg^ MHC class II^hi^ CD11c^hi^ Sirpα^neg/pos^. **(C)** On day 7, BM cells were stimulated with Class A CpGs (ODN1585) overnight. IFNα levels were determined in the supernatant using ELISA and maturation markers were evaluated by flow cytometry. Conditions are shown with and without the addition of CpGs. The individual dots represent the average of technical replicates in one independent experiment, with a total of three independent experiments performed. Error bars indicate mean ± SD. *p < 0.05, **p < 0.01, ***p < 0.001, ****p < 0.0001.

To identify mature cell subsets in the spleen as described in Figure 5, splenocytes were incubated with 10 µg/mL of anti-CD16/32 (clone 2.4G2, in-house production) and Fixable Viability Dye eFluor 780 (eBioscience). Next, we stained the cells for 20 minutes at 4 °C with the following combination of antibodies: anti-XCR1 (Clone ZET; Biolegend), anti-Ly6C (Clone HK1.4; Biolegend), anti-MCHII (Clone M5/114.15.2; Biolegend), anti-CX3CR1 (Clone SA011F11; Biolegend), anti-CD11c (Clone N418; Biolegend), anti-BST2 (Clone 927; Biolegend), anti-CD11b (Clone M1/70; Biolegend), anti-CD3 (Clone 145-2C11; Immunotools), anti-NK1.1 (Clone PK136; Immunotools), anti-B220 (Clone RA3-6B2; Biolegend), anti-Ly6G (Clone 1A8; Biolegend), anti-F4/80 (Clone BM8; Biolegend), anti-CD19 (Clone 6D5; Biolegend), anti-Siglec-H (Clone REA819; Miltenyi Biotec) and anti-Sirpα (Clone P84; Biolegend). Finally, the cells were fixed using 2 % PFA (Electron Microscopy Sciences) and subsequently acquired on the Aurora spectral flow cytometer (Cytek).

**Figure 5.**
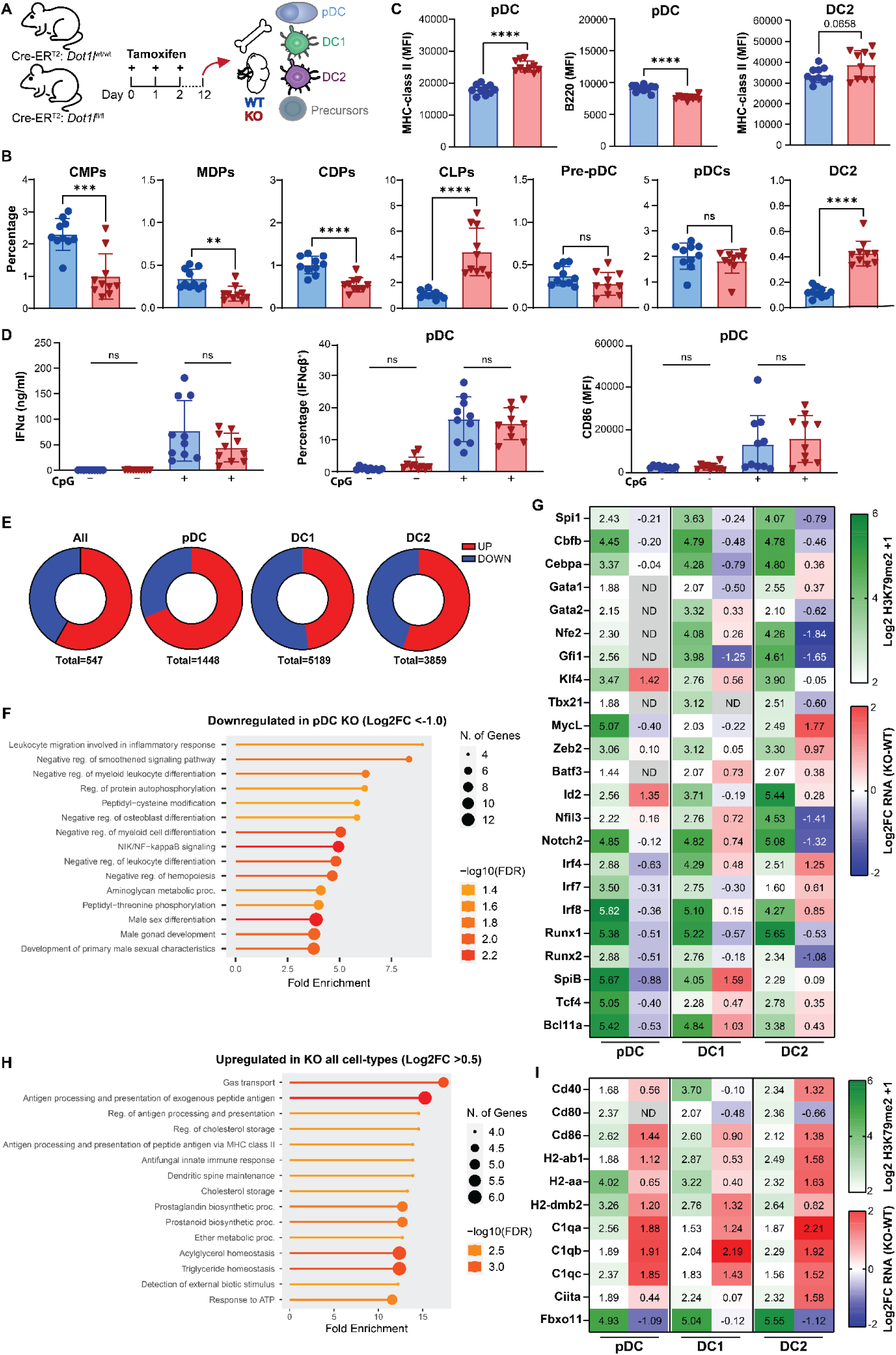
In vivo deletion of Dot1l affects specific myeloid and lymphoid precursor subsets. **(A)** Schematic overview of the experimental setup. Cre-ER^T2^ *Dot1l*^wt/wt^ and Cre-ER^T2^ *Dot1l*^fl/fl^ mice were injected with tamoxifen on three subsequent days to induce KO of *Dot1l*. On day 12, BM was harvested and analyzed. **(B)** Comparison of myeloid and lymphoid precursor percentages of Live Lineage^neg^ AF^neg^ cells in *Dot1l* WT (Cre-ER^T2^ *Dot1l*^wt/wt^) and *Dot1l* KO (Cre-ER^T2^ *Dot1l*^fl/fl^) conditions. CMPs, MDPs and CDPs were gated from MHC class II^neg^ CD11c^neg^ CD135^pos^ cells and defined as CD117^hi^ CD115^neg^ CMPs, CD117^int^ CD115^hi^ MDPs and CD117^neg^ CD115^hi^ CDPs. CLPs were defined as MHC class II^neg^ CD11c^neg^ CD135^neg^ CD127^pos^. pre-pDCs were gated as CD117^neg^ CD135^pos^ CD11c^pos^ Siglec-H^pos^ Ly-6D^pos^, pDCs as B220^pos^ Siglec-H^pos^ BST2^pos^ and cDC2s as B220^neg^ MHC class II^hi^ CD11c^hi^ Sirpα^pos^. **(C)** Comparisons of specific markers on pDCs and DC2s that were differentially expressed between *Dot1l* WT and *Dot1l* KO conditions based on the MFI of (N=10) biological replicates. **(D)** On day 12, BM cells were stimulated with Class A CpGs (ODN1585). IFNα levels were determined in the supernatant using ELISA. Additionally, cells were stained intracellularly for IFNαβ. After overnight stimulation, cells were also stained for maturation markers. Conditions are shown with and without the addition of CpGs. The individual dots represent individual mice with a total of two independent experiments performed. **(E)** Pie-charts highlighting the number of up- and down-regulated genes in *Dot1l-KO* DC populations compared to *Dot1l-WT* as determined by RNAseq (based on Log2FC <-0.5 and >0.5 for the leftmost plot and <-1.0 and >1.0 for all others). **(F)** GO-enrichment of genes that were downregulated in KO pDCs (Log2FC <-1.0). **(G)** Heatmap of selected transcription factors showing H3K79me2 signal near the transcriptional start site (TSS; N=2, green gradient) and expression differences between Dot1l-KO and WT (N-1, blue-red gradient) in the sorted DC subsets. **(H)** GO-enrichment of genes that were upregulated in expression in all sorted subsets (Log2FC >0.5). For (F) and (H) pathways were ranked by fold-enrichment and visualized using ShinyGO (*73*). **(I)** Heatmap correlating H3K79me2 TSS signal to expression differences for genes from pathways enriched in all KO subsets (as visualized in **(H)**). Error bars indicate mean ± SD. *p < 0.05, **p < 0.01, ***p < 0.001, ****p < 0.0001.

### Class A CpG stimulation of single cell suspensions

Where indicated, BM cells and/or splenocytes were stimulated with Class A CpGs (ODN1585; Invivogen, San Diego, CA, USA). 3×10^6^ BM cells or splenocytes were seeded in a round-bottom 96-wells plate with medium (RPMI-1640 (Gibco) with 10 % heat-inactivated FCS (Biowest), 50 U/mL penicillin (Lonza), 50 mg/mL streptomycin (Lonza) and 50 µM 2-mercaptoethanol) containing 1 μM ODN1585 (InvivoGen). After overnight incubation, the supernatant was harvested for ELISA, and the cells were stained with the previously described panels plus anti-CD86 (Clone GL1; BD Biosciences).

To analyze the IFNα production in the supernatant of these cultures, MaxiSorp ELISA plates (NUNC) were coated in coating buffer (pH 9.2) with 200x diluted anti-IFNα1 capture antibody (Biolegend) overnight. After, the plates were washed 3 times with PBS containing 0.05 % Tween-20. Next, the wells were blocked with 1 % PBS-BSA for 30 min at 37 °C. 100x (BM samples) or 5x (spleen samples) diluted supernatant or IFNα1 standard (Biolegend) was added to the appropriate wells and incubated for 2h on a shaking plate at RT. The plates were washed 4 times with PBS-Tween20 and subsequently incubated with 200x diluted anti-IFNα1 detection antibody (Biolegend) for 1h at RT. The plates were washed 4 times with PBS-Tween20 and incubated with 1000x diluted Avidin-HRP (Biolegend). After washing again for 5 times with PBS-Tween20, TMB (100 μg/ml) was added as a substrate and absorbance was measured at 450 nm with a microplate absorbance spectrophotometer (BioRad).

To determine the intracellular production of IFNαβ, 3×10^6^ BM cells or splenocytes were seeded in a round-bottom 96-wells plate with medium (RPMI-1640 (Gibco, Life Technologies) with 10 % heat-inactivated FCS (Biowest), 50 U/mL penicillin (Lonza), 50 mg/mL streptomycin (Lonza) and 50 µM 2-mercaptoethanol) containing 1 μM of ODN1585 (InvivoGen). After 3h incubation, Golgiplug (BD Biosciences) was added for an additional 2h. The cells were then stained extracellularly with the panels as described above and fixed using 2 % PFA (Electron Microscopy Sciences). After 2 washes with 0.5% saponin, an intracellular staining was performed with a combination of anti-IFNα (Clone RMMA-1; R&D Systems, Minneapolis, MN, USA; labeled in-house with AF647) and anti-IFNβ (Clone RMMB-1; R&D Systems; labeled in-house with AF647) in 0.5% saponin for 30 minutes at 4 °C.

### Cell sorting, RNA-sequencing preparation and data processing

*Dot1l* was genetically ablated *in vivo* via tamoxifen injections as described above. On day 12 after the first tamoxifen injection, BM cells were isolated as described above and pooled per group to prepare for sorting (*Dot1l* WT versus *Dot1l* KO). The cells were resuspended in MACS buffer (PBS, 0.5% BSA and 2 mM EDTA) and incubated with a mixture of CD11c and BST2 MicroBeads UltraPure (Miltenyi Biotec; 100 μL per 10^8^ cells) for 15 minutes at 4 °C for positive selection of CD11c^+^ and BST2^+^ cells. The cells were then washed with MACS buffer and loaded onto a MACS LS column (Miltenyi Biotec) in a MACS Separator according to the instructions of the manufacturer. The selected cells were incubated with 10 µg/mL of anti-CD16/32 (clone 2.4G2, in-house production) and Fixable Viability Dye eFluor 780 (eBioscience). Next, we stained the cells for 20 minutes at 4 °C with the following combination of antibodies: anti-XCR1 (Clone ZET; Biolegend), anti-CD11c (Clone N418; Biolegend), anti-BST2 (Clone 927; Biolegend), anti-Siglec-H (Clone 440c; BD Biosciences) and anti-Sirpα (Clone P84; BD Biosciences). Immediately after this, cDC1s, cDC2s and pDCs were sorted from the *Dot1l* WT as well as the *Dot1l* KO group using the BD FACSAria™ Fusion Flow Cytometer (BD Biosciences). pDCs were sorted as CD11c^+^ BST2^+^ Siglec-H^+^, cDC1s were sorted as CD11c^+^ BST2^-^ Siglec-H^-^ XCR1^+^ and cDC2s were sorted as CD11c^+^ BST2^-^ Siglec-H^-^ Sirpα^+^.

Sorted cells were pelleted at 1000 RCF for 5 minutes, washed once with 1 mL ice-cold PBS, transferred to RNAse/DNAse-free Eppendorf tubes and pelleted once more (1000 RCF, 4°C, 10 minutes). Pellets were re-suspended in RLT buffer (Qiagen) supplemented 1:100 with 2-mercaptoethanol (14.3M, Sigma Aldrich) and stored at -80°C. Total RNA was isolated using the RNeasy Mini kit (Qiagen) according to manufacturer’s protocol. Quality control was performed on the 2100 Bioanalyzer using the RNA nanochip (Agilent), and samples were library amplified using the SMART-Seq® v4 Ultra® Low Input RNA Kit (Takara) and quality checked on the 2100 Bioanalyzer using the DNA 7500 chip (Agilent, Santa Clara, CA, USA). Samples were sequenced (51bp, paired end) using the NovaSeq 6000. Reads were aligned against the mouse reference genome (GRCm38/mm10) using Tophat (version 2.1, bowtie version 1.1), supplied with a Gene Transfer File (GTF, Ensembl version 77) using the following parameters: “prefilter-multihits –no-coverage-search – bowtie1 –library-type fr-firststrand”. A custom script based on HTSeq-count was used to count the number of uniquely mapped reads per gene, as listed in the provided Gene Transfer Format (GTF) file.

Subsequent analysis was performed in R version 4.2.2 with Bioconductor packages from release 3.16. The analysis was restricted to the genes that had at least 10 counts 3 or more samples to exclude very low abundance genes. On these genes, a principal component analysis (PCA) was performed using the “prcomp” function on variance stabilizing transformed data from the “vst” function of the DESeq2 package (version 1.38.3). Data was visualized using ggplot2 (version 3.5.1). GO enrichment analyses were performed and visualized using ShinyGO v0.82, hosted by the South Dakota State University, by filtering for a minimum pathway size of 20 and ranking based on fold enrichment but otherwise using default settings (*71–73*). For these analyses, only genes that had an expression of at least 2^5^ counts per million reads in either WT or KO as well as a Log2FC of >1 or <-1, (unless otherwise indicated) were considered. To facilitate quantitative comparisons with the H3K79me2 ChIPseq data (see below) the mean library-size corrected expression value for each gene between all samples (BaseMean) was calculated and, where relevant (for Figure 1G) further normalized for transcript length using the ‘fpkm‘ function of DESeq2 (version 1.38.3).

### Normalized H3K79me2 ChIP-seq preparation and data processing

BM cells from WT C57BL/6J mice were sorted as described above into pDC, cDC1 and cDC2 pools, three pools each with each pool consisting of cells from 5 mice. Sorted cells were cross-linked for ten minutes at room temperature in RPMI supplemented with 1 % methanol free formaldehyde (Thermo Scientific) and subsequently quenched by adding glycine (final concentration 125 mM) for five minutes. Samples were then washed twice using PBS + protease inhibitor cocktail (PIC; Roche) + 10% FBS (Capricorn), aspirated, and re-suspended in ice-cold nuclei lysis buffer (50 mM Tris-HCl pH8, 10 mM EDTA pH8, 1 % SDS + PIC) for 10 minutes. Samples were sonicated for three minutes (3x 30sec on, 30sec off) using the BioRuptor (Pico), after which debris was pelleted (4°C, 13000 RCF, 10 minutes) and supernatant transferred to a new Eppendorf tube. Samples were supplemented with sample buffer (9 parts ChIP dilution buffer (50 mM Tris-HCl pH 8, 0.167 M NaCl, 1.1 % Triton X-100, 0.11 % sodium deoxycholate) and 5 parts RIPA-150 (50 mM Tris-HCl pH8, 0.15M NaCl, 1 mM EDTA pH8, 0.1 % SDS, 1 % Triton X-100, 0.1 % sodium deoxycholate + PIC) per 1 part sample. Ten percent of each suspension de-crosslinked by adding 10 mg/mL RNAse A (Sigma Aldrich) and 10 mg/mL Proteinase K (Sigma Aldrich) and incubating for 1 hour at 50°C, then overnight at 65°C. From these de-cross-linked samples, the shearing efficacy was checked by gel electrophoresis, and gDNA was isolated and quantified (as described above).

A yeast chromatin spike-in was added 1:1000 to all samples based on DNA concentration. This spike-in was generated using a *dot1Δ*, H3-3xHA (HHT2-3xHA) yeast strain prepared as described previously (*74*). Samples were homogenized in sample buffer, and 10% was stored as input while 90% was used for simultaneous IP for HA (yeast spike-in) and H3K79me2, using 0.5 μg mouse anti-HA antibody (clone 12CA5, Protein Facility NKI) and 2 µl rabbit anti-H3K79me2 (Millipore) per sample. Samples were incubated overnight under rotation at 4°C, after which Dynabeads™ Protein G (Thermo Scientific) in sample buffer were added for 2 hours. Supernatant was aspirated using the DynaMag-2 magnetic rack (Thermo Scientific), and samples were washed one with RIPA-150 (50mM Tris-HCl pH8, 0.15M NaCl, 1mM EDTA pH8, 0.1% SDS, 1% Triton X-100, 0.1% sodium deoxycholate) twice with RIPA-500 (50 mM Tris-HCl pH 8, 0.5 M NaCl, 1 mM EDTA pH8, 0.1 % SDS, 1 % Triton X-100, 0.1 % sodium deoxycholate), twice with RIPA-LiCl (50 mM Tris-HCl pH 8, 1 mM EDTA pH 8, 1 % Nonidet P-40, 0.7 % sodium deoxycholate, 0.5 M LiCl2) and once with TE. Samples were re-suspended in 150 μL Direct Elution buffer (10 mM Tris-HCl pH 8, 0.3 M NaCl, 5 mM EDTA pH8, 0.5 % SDS) de-crosslinked and quantified as described above. Samples were library amplified, pooled, and sequenced (51bp, paired end) on the NovaSeq 6000. Reads were aligned to a concatenated mouse (GRCm38/mm10) and yeast (sacCer3) genome, on which peaks were called using MACS2 version 2.2.9.1 with the arguments ‘-f BEDPE –g mm –keep-dup all --broad’. Data was further normalized using the spike-in reads and visualized as described previously (*74*).

PCA was performed on the ChIPseq signal near the transcriptional start site (TSS; + 2kb) using the “prcomp” function as described above. Peaks that uniquely mapped to a single sorted DC subtype were further analyzed with the PubMed enrichment built into the STRING database using default arguments (*75*). Heatmaps and correlation plots with H3K79me2 signal near the TSS were visualized using GraphPad Prism (version 10.0.2).

### Analysis of flow cytometry data

All data was unmixed after acquisition using SpectroFlo software (Cytek). Further analysis was performed using OMIQ by Dotmatics. Manual gating was used to identify cell subsets after removing debris and gating on live cells. Data was generated on the geometric mean fluorescent intensity (gMFI) of specific markers or percentage change of target populations, and this was plotted in GraphPad Prism v8 (GraphPad).

### Statistical analysis

To identify statistical differences between 2 groups, a two-tailed t-test was used. For the DOT1L inhibitor experiments, a paired two-tailed t-test was used. For 3 or more groups, a one-way or two-way ANOVA with a Šidák post-hoc test was performed. All statistical analyses were performed in GraphPad Prism v8 (GraphPad).

## Results

### Dot1l expression and H3K79me2 methylation state in DC subsets

To explore the potential importance of DOT1L in the development of DCs, we first investigated the presence of *Dot1l* mRNA in DC (precursors) through publicly available RNA sequencing datasets. A scRNA sequencing dataset from Schlitzer *et al.* (2015) showed that DC progenitors from the BM indeed express *Dot1l* (Figure 1A) (*13*). Moreover, bulk RNA sequencing data from sorted mature DCs in the spleen from Rodrigues et al. (2023) revealed the presence of *Dot1l* in cDC1s, cDC2s and pDCs, although with some variation between samples (Figure 1B) (*17*).

Since *Dot1l* appeared to be expressed in several DC subtypes, we sorted pDCs (Siglec-H^pos^ BST2^pos^), cDC1s (CD11c^pos^ XCR1^pos^), and cDC2s (CD11c^pos^ Sirpα^pos^) from WT C57BL/6 mice and performed ChIPseq for H3K79me2 as well as RNAseq. While DOT1L is responsible for depositing H3K79me1 and H3K79me2 in transcribed regions, H3K79me2 shows a stronger correlation with gene expression and has been studied more extensively. To quantitatively compare H3K79me2 peaks and levels between the DC populations, a yeast chromatin spike-in was added to all samples in parallel and co-immuno-precipitated as recently described (Figure 1C) (*74*). Within actively transcribed genes, H3K79me2 was predominantly deposited between the transcriptional start site (TSS) and the first internal exon (FIE), consistent with previous findings in other cell types (Figure 1D) (*39,41,53*). Comparison of the spike-in normalized peaks revealed that global H3K79 methylation levels were comparable between the sorted DC populations (Supplementary Figure S1A), indicating that DOT1L has similar activity in the different DC subsets.

Analysis of the methylated regions revealed that some genes had similar H3K79me2 levels in all subtypes (*e.g. Runx1*, Figure 1E, top), while others were only methylated in one DC population (*e.g. Tcf4*, Figure 1E, bottom), Overall, 4522 H3K79me2 peaks were identified as shared between all sorted cell populations, while 2161 peaks uniquely mapped to pDC, 3096 to cDC1 and 2647 to the cDC2 population (Figure 1F; Supplementary Figure S1B). GO analyses revealed that the pDC-specific peaks were consistent with gene signatures from previous pDC related publications, suggesting that *Dot1l* might indeed be involved in the regulation of key genes within this cell population (Supplementary Figure S1C).

As H3K79me2 methylation typically correlates with active transcription (*51,52,55,76*), we performed RNAseq to confirm whether differences in H3K79me2 deposition between the DC populations related to transcriptional differences. To compare the ChIPseq and RNAseq data, we quantified the H3K79me2 signal near the TSS (Supplementary Figure S1D) and correlated this to transcript length normalized gene expression (Fragments Per Kilobase of exon per Million mapped fragments – FPKM). Generally, the H3K79me2 signal near the TSS correlated moderately with transcription. The correlation was strongest in pDCs (ρ = 0.53), followed by cDC2s (ρ = 0.46) and weakest in the cDC1 (ρ = 0.38) population (Figure 1G). For some genes encoding transcription factors that regulate DC differentiation and function (*Tcf4, Bcl11a, Irf8 and SpiB*) high levels of both H3K79me2 and mRNA expression were observed in the sorted pDCs, suggesting that H3K79me2 was linked to transcriptional activity (Figure 1H). In conclusion, the normalized ChIPseq data indicate that DOT1L induces similar levels of global H3K79me2 in DC subsets but also show specific differences in specific genes in certain DC subsets. As these changes are, in part, accompanied by changes in expression, this suggests that DOT1L mediated H3K79me2might be involved in directly or indirectly regulating DC differentiation, especially for the pDC subset.

### Decreased development of pDCs and increased development of cDC2s upon in vitro deletion of Dot1l in FLT3L- and SCF-supplemented BM cultures

Considering that the ChIPseq revealed that *Dot1l* is active in DCs and methylated common and unique genes, we next investigated what role *Dot1l* plays in DC development using an *in vitro* FLT3L/SCF BM DC culture system (*77,78*). We used a mouse strain (Cre-ER^T2^;*Dot1l*^fl/fl^) in which the genetic ablation of *Dot1l* (KO; loss of exon 2) can be induced through the addition of 4-hydroxytamoxifen (4-OHT) (Figure 2A). It should be noted that after 4-OHT-induced *Dot1l*-KO, the loss of DOT1L-associated H3K79 methylation is delayed as it primarily depends on dilution by cell division (*79*). FLT3L/SCF-supplemented Cre-ER^T2^;*Dot1l*^wt/wt^ (WT) BM culture led to the generation of BST2^pos^ Siglec-H^pos^ pDCs and CD11c^pos^ MHC class II^pos^ Sirpα^pos^ cDC2s as well as CD11c^pos^ MHC class II^pos^ XCR1^pos^ cDC1s on day 7 (Figure 2B-C; Supplementary Figure S2). All subsets were pre-gated as Live CD45^pos^ Lineage^neg^ cells. Loss of *Dot1l* through *in vitro* treatment with 4-OHT decreased the number and frequency of pDCs from 15.4% to 4.2% on day 7 after the start of the culture (percentage of live CD45^pos^ Lineage^neg^ cells; Figure 2C-D; Supplementary Figure S2). In contrast, the number and frequency of cDC2s increased from 9.7% to 16.9% in *Dot1l*-KO cultures while cDC1s were unaffected (Figure 2C-D; Supplementary Figure S2). In conclusion, KO of *Dot1l* in total murine BM decreased the number of pDCs and increased the number of cDC2, after culturing for 7 days with FLT3L and SCF.

### In vivo deletion of Dot1l in the bone marrow followed by in vitro expansion leads to reduced pDCs and myeloid precursors

Deletion of *Dot1l in vitro* appeared to affect the development of DCs in BM cultures. While DOT1L has a short half-life of approximately 2 hours (*80,81*), H3K79me is a stable histone modification and loss of the epigenetic mark upon inactivation of DOT1L has been reported to be mainly dependent on dilution by cell division (*79,82*). Proliferation during the 7-day BM cultures likely facilitated the loss of H3K79me following the loss of DOT1L in an early stage of the cultures. To explore whether *in vivo* loss of DOT1L protein led to a direct effect on DC (precursor) frequency, we treated the Cre-ER^T2^;*Dot1l*^fl/fl^ and Cre-ER^T2^;*Dot1l*^wt/wt^ mice *in vivo* with tamoxifen to generate *Dot1l*-deficient mice. The mice were injected intra-peritoneally with tamoxifen on three consecutive days after which the genetic ablation of *Dot1l* was confirmed by PCR (Supplementary Figure S3A-C). On day 3, cells isolated from the spleen and BM were stained with an antibody panel to identify DC precursors and mature cell subsets. The identification of myeloid and lymphoid precursors in the BM was based on the presence or absence of previously described markers on these cell subsets (Supplementary Figure S3A) (*1,22,83–87*). All subsets were pre-gated as Live, Lineage^neg^ Autofluorescence (AF)^neg^ cells. Subsequently, CMPs, MDPs and CDPs were gated from MHC class II^neg^ CD11c^neg^ CD135^pos^ cells and defined as CD117^hi^ CD115^neg^ CMPs, CD117^int^ CD115^hi^ MDPs and CD117^neg^ CD115^hi^ CDPs. CLPs were defined as MHC class II^neg^ CD11c^neg^ CD135^neg^ CD127^pos^, pre-pDCs were gated as CD117^neg^ CD135^pos^ CD11c^pos^ Siglec-H^pos^ Ly-6D^pos^, pDCs as B220^pos^ Siglec-H^pos^ BST2^pos^ and cDC1s and cDC2s as B220^neg^ MHC class II^hi^ CD11c^hi^ Sirpα^neg/pos^ (Supplementary Figure S3A). On day 3 we did not observe changes in precursor or mature cell populations in the BM (Supplementary Figure S3D) or the spleen (Supplementary Figure S3E-F). These results indicate that loss of DOT1L protein itself does not result in direct changes in DC subsets and precursors and suggests that more extensive proliferation is necessary to lose H3K79me. Moreover, if the effect of DOT1L is at the level of bone marrow precursors then effects on mature DC subsets will be delayed as differentiation of mature DC subsets will require 5-7 days (*88*).

To recapitulate our previous results and investigate changes in DC precursors in a BM culture setting, isolated BM cells from tamoxifen-treated Cre-ER^T2^;*Dot1l*^fl/fl^ and Cre-ER^T2^;*Dot1l*^wt/wt^ mice were cultured with FLT3L and SCF as described above (Figure 3A). On day 7 the cultures were harvested and stained to identify precursors and mature cell subsets (Supplementary Figure S3A). In line with the previous *in vitro* BM cultures, pre-pDCs as well as pDCs showed a clear reduction, while cDC2s were increased upon *Dot1l* KO (Figure 3B). A clear reduction in the percentage of cells belonging to the myeloid linage was observed, while the lymphoid compartment and the cDC1 subset remained unaffected in the *Dot1l*-KO cultures (Figure 3B). CMPs, MDPs and CDPs were all significantly reduced after *Dot1l* KO in contrast to CLPs, which remained unaffected.

pDCs have been described to respond to class A CpGs by rapidly producing large amounts of type I IFNs (*10,12*). A recent paper additionally observed type I IFN production by pre-cDC2s after class A CpG stimulation (*89*). To determine whether the remaining DCs from *Dot1l*-deficient BM cultures were still functional, BM cells were stimulated with class A CpGs (ODN 1585) overnight on day 7 and IFNα levels in the supernatant were detected using ELISA. This revealed that IFNα production after CpG stimulation was absent in the *Dot1l-*KO cultures (Figure 3C), which correlates with the decreased number of pDCs. However, the remaining *Dot1l*-KO DCs still upregulated the maturation marker CD86 in response to class A CpGs (Figure 3D). To summarize, *in vivo* deletion of *Dot1l* did not result in changes in immune populations in the BM and spleen on day 3 after the first tamoxifen injection. However, a subsequent 7-day culture of the BM cells resulted in a reduction of myeloid precursors, pre-pDCs and pDCs and upregulation of cDC2s. This indicates that loss of DOT1L alone is not sufficient to observe the changes in DC subsets and that proliferation and/or differentiation is necessary.

### In vitro inhibition of DOT1L in WT BM cultures results in decreased generation of pDCs and increased generation of cDC2s

DOT1L is the sole methyltransferase of H3K79, but has also been reported to have cellular roles independent of H3K79 methylation (*63–67,90*). To investigate the role of the catalytic activity of DOT1L in DC differentiation, we treated the cells in our culture system with the potent and selective DOT1L inhibitor SGC-0946 (Figure 4A) (*57*). Treatment with SGC-0946 did not impact the viable cell count of the FLT3L/SCF-BM cultures (Supplementary Figure S4). Similar to the *Dot1l* KO cultures, we observed a decrease in CDPs, pre-pDCs and pDCs after 7 days culture in the presence of SGC-0946. Furthermore, cDC2s were increased and the fraction of CLPs was unaffected. However, no changes were observed in CMP and MDP populations (Figure 4B). In line with our previous results, the ability of CpG-stimulated DCs to produce IFNα was completely abolished after DOT1L inhibition, while CD86 expression did not change significantly (Figure 4C). In short, inhibition of the methyltransferase activity of DOT1L resulted in similar effects as *Dot1l* KO. Since DOT1L inhibitor treatment does not affect the stability of the DOT1L protein (*91*), this indicates that the phenotype induced by DOT1L is mainly due to its catalytic activity.

### In vivo deletion of Dot1l affects specific myeloid and lymphoid precursor subsets

Although the *in vivo* genetic ablation of *Dot1l* in Cre-ER^T2^;*Dot1l*^fl/fl^ mice did not result in changes in the DC (precursor) populations 3 days after the first treatment with tamoxifen, the subsequent 7-day *in vitro* culture altered DC precursor and mature DC subset frequencies. Combined with the DOT1L inhibitor experiments, this supports the hypothesis that the effect of DOT1L on DC differentiation is associated with its methyltransferase activity. Earlier studies estimated the half-life of DCs to be 5-7 days (*88*). Therefore, to accommodate for both the time needed to passively dilute H3K79me2, as well as for subsequent differentiation, we set up an *in vivo* experiment in which we evaluated the cell populations in the spleen and BM on day 12 after the start of tamoxifen administration (Figure 5A). *Dot1l* deletion efficiency was more than 90% in both BM and spleen (Supplementary Figure S5A), while staining for H3K79me2 revealed that approximately 80% of BM cells and 10% of splenocytes showed reduced H3K79me2 levels (Supplementary Figure S5B-C). Unfortunately, it was not possible to determine H3K79me2 staining in specific cell subsets, as the harsh denaturation conditions required for this staining could not be combined with the elaborate panel of surface markers needed for the identification of these cells.

On day 12 after the first dose of tamoxifen, we observed a decrease in CMPs, MDPs and CDPs, and an increase in CLPs as well as cDC2s in the BM, similar to the *in vitro* BM cultures (Figure 5B). However, no significant difference was observed in the frequency of pre-pDC and pDC populations (Figure 5B). Flow cytometry analyses revealed a significant increase in MHC class II expression on pDCs and to a lower degree on cDC2s and a decrease in B220 expression on pDCs in *Dot1l*-KO mice compared to *Dot1l*-WT mice (Figure 5C). BM *Dot1l*-KO pDCs retained the ability to upregulate CD86 and produce type I IFNs after stimulation with class A CpGs as detected by intracellular IFNαβ staining (Figure 5D). In the spleen, no differences in the frequencies of cDC1s, cDC2s and pDCs subsets were observed (Supplementary Figure S5D), although a strong increase in CD11b expression on pDCs, cDC1s and cDC2s as well as an increase in Ly6C expression on pDCs was detected in *Dot1l*-KO mice (Supplementary Figure S5E). After overnight stimulation of splenocytes with class A CpGs, *Dot1l*-KO pDCs still produced type I IFNs and upregulated maturation marker CD86 (Supplementary Figure S5F).

In summary, after 12 days of *Dot1l* deletion *in vivo* we detected changes in DC precursor and cDC2 frequencies in the BM, but not in the spleen. In addition, phenotypic changes were observed in DC subsets from both organs. In contrast to the *in vitro* experiments, pDCs were not reduced in *Dot1l*-KO mice and retained their ability to produce type I IFNs and to upregulate maturation marker CD86.

To gain further insight into the consequences of *Dot1L* loss in vivo, we performed RNAseq on sorted WT and KO populations (Siglec-H^pos^ BST2^pos^ pDC, CD11c^pos^ XCR1^pos^ cDC1 and CD11c^pos^ Sirpα^pos^ cDC2) from the BM at day 12 after the start of tamoxifen treatment. Principal component analysis revealed that KO cells clustered closest to their WT counterparts, suggesting that the global changes upon *Dot1l* loss were modest (Supplementary Figure S5G,). Overall, 228 genes were downregulated in expression in all KO cell-types (Log2FC <-0.5) and based on GO analysis these genes were mostly related to metabolic pathways and the regulation of transcription (Figure 5E, Supplementary Figure S5H). In contrast, genes that were downregulated in sorted *Dot1l-*KO pDC (452 genes based on Log2FC <-1.0) enriched for several pathways that negatively regulate (myeloid) cell differentiation, perhaps underlying the loss of pDC identity as previously observed upon subsequent cell culture *in vitro* (Figure 5F). In line with this, several transcription factors involved in pDC differentiation (*e.g. Runx2, Tcf4, Bcl11a, Irf8 and SpiB*) were modestly downregulated in *Dot1L-*KO pDC, while changes for the remaining transcription factors and cell-types were more variable (Figure 5G).

Similarly, most upregulated genes were unique in each cell type, with minimal overlap between the cell types. Nevertheless, genes upregulated in all KO populations (319 genes based on Log2FC >0.5) enriched for various antigen presentation and activation related pathways (Figure 5H). This corroborates earlier results (*e.g.* Figure 5C) that suggest a more pro-inflammatory state in *Dot1l*-KO BM and spleen. Correlating transcriptional changes in KO with H3K79me2 signal (in WT) revealed that the genes underlying these upregulated signatures were generally hypo-methylated (Figure 5I). This suggests that these genes are likely indirect targets of the loss of H3K79me2. Among the enriched activation-related genes were several activation markers (*Cd40*, *Cd80*, *Cd86*), classical complement factors (*C1qa*, *C1qb*, *C1qc*) and MHC class II complex members (*H2-ab1*, *H2-aa*, *H2-dmb2*). Expression of the latter is predominantly mediated through the master regulator CIITA and its negative regulator FBXO11 (*92*). *Ciita* was H3K79 hypo-methylated in all cell-types, while *Fbxo11* was hyper-methylated (Figure 5I), suggesting that *Dot1l* might regulate MHC class II expression levels indirectly via *Fbxo11* (Figure 5I, Supplementary Figure S5I). *Dot1l-*KO pDCs and cDC2s showed an increase in *Ciita* expression and a decrease in *Fbxo11* expression, which might underlie the increased antigen presentation signatures predominantly observed in these two cell types (Figure 5I). Lastly, while HDACs have been linked to both DC differentiation and *Dot1l* function, we observed no consistent changes in their expression upon *Dot1l*-KO, suggesting that the potential interplay between these proteins is not mediated solely through H3K79me2 levels (Supplementary Figure S5J).

Combined, *Dot1l* potentially affects the differentiation of pDCs via positive regulation of a number of transcription factors, while also suppressing the basal activation state of all the sorted DC subtypes. Further studies will be required to further elucidate the mechanism underlying the regulation of these differentiation and activation related pathways.

## Discussion

DCs serve as key regulators of the overall immune response. Their differentiation trajectory is complex, involving both lymphoid and myeloid lineages. The role of epigenetic mechanisms guiding DC differentiation is still relatively unexplored (*1,29,30*). Recently, histone deacetylases and a demethylase have been implicated in the differentiation of various DC subtypes (*33–36*) but otherwise, the role of epigenetic mechanisms in DC development remains understudied. Within this study, we identified a role for the histone methyltransferase DOT1L in DC differentiation towards the pDC, cDC1 and cDC2 subtypes. Previous studies have described a central role for methyltransferase DOT1L in the differentiation of other immune cell types, such as macrophages, NK cells, B and T lymphocytes cells (*39–42,44–46*), and recently DOT1L has been linked to DC function as well (*48,49*). However, to our knowledge, the impact of DOT1L on the differentiation of specific DC progenitors towards mature DC subsets remains largely unexplored.

DOT1L is a histone methyltransferase that specifically deposits H3K79me on active genes. H3K79me2 ChIPseq on sorted DC populations revealed a distribution pattern that started around the TSS and dropped off after the first internal exon was reached, a pattern consistent with that observed in other cell types (*39,41,53,74*). While there was substantial overlap in H3K79me2 between the three subsets, pDCs, cDC1s and cDC2s all contained distinct peaks as well, which suggests differential regulation of these subtypes by DOT1L. The correlation between H3K79me2 signal around the TSS and transcription based on RNAseq was strongest in pDCs. Intriguingly, several transcription factors involved in pDC development, including *Irf8, SpiB, Tcf4,* and *Bcl11a,* were abundantly methylated in pDCs. This observation pointed to a potential role for DOT1L in the regulation of DC differentiation and especially of pDC.

When DOT1L was genetically ablated *in vitro*, we observed a decrease in pDC differentiation, while cDC2 numbers increased and cDC1s were unaltered after 7 days culture with FLT3L and SCF. *In vivo* KO of *Dot1l* did not result in changes in DCs or their precursors in the BM on day 3, but a subsequent 7-day culture period again revealed again changes in pDC and DC2 populations, as well as a decrease in myeloid precursors. These DC cultures were unable to produce IFNα upon Class A CpG stimulation which was in line with the decreased pDC frequencies. *Ex vivo* analysis of *in vivo Dot1L* KO BM after 12 days demonstrated a decrease of myeloid precursors and an increase of cDC2s and lymphoid precursors, but unexpectedly no changes in the frequency of pDCs. However, analysis of sorted pDCs at this time point revealed that the cells had changed their phenotype in the absence of DOT1L. Pathways related to *‘negative regulation of myeloid differentiation’* were specifically downregulated based on GO enrichment in *Dot1L*KO pDCs. In addition, the highly methylated transcription factors important for pDC differentiation, as *Irf8, SpiB, Tcf4,* and *Bcl11a,* showed a decrease of expression in the *Dot1L*KO pDCs. Very recent studies describe a close relationship between preDC2A and pDCs and show that preDC2a express pDC transcription factors and markers such as Tcf4, BST2 and Siglec-H and also produces IFN upon class A CpG stimulation (*89*). While we included B220 in the pDC gating in the majority of the experiments, including the detection of intracellular IFNαβ, and thereby selected for genuine pDCs and not preDC2A, this was not included in the sorting for the results shown in Figure 1 and Figure 5. Therefore, we cannot exclude that there was a contamination of preDC2A in the pDC gate in our ChIPseq and RNAseq experiments. Our observation that loss of DOT1L affects both pDC and DC2 differentiation is interesting considering the recently described shared features (*89*), but future studies will be necessary to more precisely determine the role of DOT1L in the specific differentiation trajectories of the cDC2A and B subsets as well as DC3 and other myeloid cell types.

While DOT1L mainly acts as a histone methyltransferase with specificity towards H3K79me, other non-canonical functions have been described as well (*63–66,82,90,93*) and such functions may also be at play in DCs. To potentially differentiate between the two, we performed the *in vitro* expansion using the highly specific DOT1L inhibitor SGC-0946 (*57*). Supplementing our *in vitro* BM cultures with SGC-0946 generally recapitulated the results of our *Dot1l*-KO experiments and thereby confirmed that the effect of DOT1L on DC differentiation is related to its catalytic activity most likely via H3K79me. Since H3K79me is mainly lost via dilution due to proliferation, the effects of *Dot1l-*KO can only be expected after substantial number of cell divisions (*79,94–98*). A possible explanation for the discrepancy in pDC generation *in vitro* BM cultures and *in vivo* experiments is the amount of proliferation that occurs. The addition of SCF to FLT3L BM cultures stimulates proliferation of c-KIT^+^ progenitors and this may drive faster loss of H3K79me than in vivo (*77*). In addition, pDCs are proposed to have a longer half-life than cDCs in mice and this potentially could be of importance for changes in DC2 and pDC *in vitro* versus *in vivo* (*99*). Due to the harshness of the H3K79me2 staining (*39,41,76*) we could unfortunately not co-stain with the markers needed to identify specific DC subsets, but further experiments with sorted DC subsets and later time points will be necessary to demonstrate H3K79me loss and determine potential effects on pDCs *in vivo*.

Genes that were upregulated in *Dot1l-*KO conditions often contained low or no detectable H3K79me2 and are therefore likely indirect targets of DOT1L (*51,52,55*). A clear upregulation was observed in all DC subsets regarding genes involved in antigen presentation and immune response pathways. Multiple activation-related genes (*Cd40, Cd80, Cd86*), complement factors (*C1qa, C1qb, C1qc*) and MHC class II related genes (*H2-ab1, H2-aa, H2-dmb2*) were upregulated in all DC subsets. Indeed, increased surface level expression of MHC class II was observed for pDCs and to a lower extent for cDC2s. CIITA is a known regulator of MHC class II expression, and its expression is controlled by the negative regulator FBXO11 (*92,100*). *Fbxo11* was hyper-methylated for H3K79me2 in all DC subsets, while *Ciita* was relatively hypo-methylated, suggesting that DOT1L, via H3K79me2, might indirectly regulate MHC class II expression. These results are in line with previous studies on inflammation regulation by DOT1L. A study on the function of DOT1L in atherosclerotic plaque macrophages, for example, reported that DOT1L deficiency or inhibition results in macrophage hyper-activation, MHC class II upregulation, and linked DOT1L to the regulation of inflammatory processes in these macrophages (*44*). Tang *et al.* proposed that intestinal immune tolerance is stimulated via increased DOT1L expression and H3K79me2 in DCs (*49*). Additionally, in pancreatic and colon cancer mouse models, H3K79me2 controls FOXM1 expression, which is associated with the regulation of DC maturation (*48*), indicating a role for DOT1L in DC function in different models.

Previous studies investigating epigenetic regulators showed a dominant effect of HDAC1, but not HDAC2 on pDC and cDC2 development, and of HDAC3 on pDC development (*34,36*). A recent study indicated a role for DOT1L upstream of HDAC1 and positive regulation of DOT1L expression by HDAC1 (*101*). In contrast, we previously detected negative regulation of DOT1L activity by HDAC1 (*38,91*). In our data, we observed considerable H3K79 methylation of the genes encoding HDAC1-3 in all DC subsets, leaving the possibility open that DOT1L might directly regulate the expression of HDACs in DCs. However, the effect of *Dot1l* KO on HDAC expression was variable between subsets. Further research would be required to properly investigate the interaction of DOT1L with HDACs in DCs and its potential effect on their differentiation.

Overall, our study reveals a differential role for DOT1L catalytic activity in the development of myeloid precursors, cDC2, and pDC in *in vitro* and *in vivo* settings. Furthermore, *in vivo* experiments showed that *Dot1l* deficiency results in an elevated inflammatory status of DCs. These findings form the basis for more mechanistic studies investigating how DOT1L influences DC subset differentiation and function.

## Acknowledgements

This work was supported by NWO ZonMW (TOP91218024 to JH; TOP91218022 to FvL), the Dutch Cancer Society (2019-12802, 2021-2/14093, 2022-14453 to JH; NKI2018-1/11490, 2022-2 EXPL/14479 to FvL), from Health Holland TKI-PPP to JH, and institutional grants to the Netherlands Cancer Institute from the Dutch Cancer Society (KWF) and the Dutch Ministry of Health, Welfare and Sports. AJA and JS were supported by NWO Veni ZonMw (09150162010163) and Stichting Cancer Center Amsterdam (CCA 2022-9-83). AZW was supported by Chinese Scholarship Council. We would like to thank the staff of Netherlands Cancer Institute Preclinical Intervention Unit of the Mouse Clinic for Cancer and Ageing (MCCA) for the animal care and technical support. We acknowledge the Microscopy and Cytometry Core Facility (MCCF) at the Amsterdam UMC (Location VUmc) and the Flow Cytometry facility of the NKI for providing assistance and advice with flow cytometry experiments. We would like to thank the Genome Core Facility (GCF) of the NKI for technical support for the RNAseq and ChIPseq experiments, and Teun van de Brand for support with subsequent analysis. We thank Heinz Jacobs for valuable discussions and input during the studies.

## Author contributions

Conceptualization: RB, WdL, MM, FvL, JdH

Formal analysis: RB, WdL, MM, AM, JvD

Funding acquisition: FvL, JdH

Investigation: RB, WdL, AZ, JS, VK, MNT, TvW, NST, AA

Methodology: RB, WdL, NP, TvW, AA

Data curation and Software: WdL, AM, JvD

Supervision: FvL, JdH

Validation: RB, WdL, MM

Visualization: RB, WdL, AM

Writing: RB, WdL, FvL, JdH

## Data and Materials Availability

All processed data are in the main text or the supplementary materials.

## References

(1) Anderson, D.A., Dutertre, C.-A., Ginhoux, F., and Murphy, K.M. (2021) Genetic models of human and mouse dendritic cell development and function. Nat. Rev. Immunol. 21, 101–115

(2) Cabeza-Cabrerizo, M., Cardoso, A., Minutti, C.M., Pereira da Costa, M., and Reis e Sousa, C. (2021) Dendritic Cells Revisited. Annu Rev Immunol. 39, 131–166

(3) Adams, N.M., Das, A., Yun, T.J., and Reizis, B. (2024) Ontogeny and Function of Plasmacytoid Dendritic Cells. Annu Rev Immunol. 42, 347–373

(4) Durai, V., and Murphy, Kenneth M. (2016) Functions of Murine Dendritic Cells. Immunity. 45, 719–736

(5) Lewis, Kanako L., Caton, Michele L., Bogunovic, M., Greter, M., Grajkowska, Lucja T., Ng, D., Klinakis, A., Charo, Israel F., Jung, S., Gommerman, Jennifer L., Ivanov, Ivaylo I., Liu, K., Merad, M., and Reizis, B. (2011) Notch2 Receptor Signaling Controls Functional Differentiation of Dendritic Cells in the Spleen and Intestine. Immunity. 35, 780–791

(6) Brown, C.C., Gudjonson, H., Pritykin, Y., Deep, D., Lavallée, V.-P., Mendoza, A., Fromme, R., Mazutis, L., Ariyan, C., Leslie, C., Pe’er, D., and Rudensky, A.Y. (2019) Transcriptional Basis of Mouse and Human Dendritic Cell Heterogeneity. Cell. 179, 846–863.e824

(7) Satpathy, A.T., Briseño, C.G., Lee, J.S., Ng, D., Manieri, N.A., Kc, W., Wu, X., Thomas, S.R., Lee, W.L., Turkoz, M., McDonald, K.G., Meredith, M.M., Song, C., Guidos, C.J., Newberry, R.D., Ouyang, W., Murphy, T.L., Stappenbeck, T.S., Gommerman, J.L., Nussenzweig, M.C., Colonna, M., Kopan, R., and Murphy, K.M. (2013) Notch2-dependent classical dendritic cells orchestrate intestinal immunity to attaching-and-effacing bacterial pathogens. Nat Immunol. 14, 937–948

(8) Briseño, C.G., Satpathy, A.T., Davidson, J.T.t., Ferris, S.T., Durai, V., Bagadia, P., O’Connor, K.W., Theisen, D.J., Murphy, T.L., and Murphy, K.M. (2018) Notch2-dependent DC2s mediate splenic germinal center responses. Proc. Natl. Acad. Sci. U.S.A. 115, 10726–10731

(9) Tussiwand, R., Everts, B., Grajales-Reyes, Gary E., Kretzer, Nicole M., Iwata, A., Bagaitkar, J., Wu, X., Wong, R., Anderson, David A., Murphy, Theresa L., Pearce, Edward J., and Murphy, Kenneth M. (2015) Klf4 Expression in Conventional Dendritic Cells Is Required for T Helper 2 Cell Responses. Immunity. 42, 916–928

(10) Reizis, B. (2019) Plasmacytoid Dendritic Cells: Development, Regulation, and Function. Immunity. 50, 37–50

(11) Ngo, C., Garrec, C., Tomasello, E., and Dalod, M. (2024) The role of plasmacytoid dendritic cells (pDCs) in immunity during viral infections and beyond. Cell. Mol. Immunol. 21, 1008–1035

(12) Swiecki, M., and Colonna, M. (2015) The multifaceted biology of plasmacytoid dendritic cells. Nat. Rev. Immunol. 15, 471–485

(13) Schlitzer, A., Sivakamasundari, V., Chen, J., Sumatoh, H.R.B., Schreuder, J., Lum, J., Malleret, B., Zhang, S., Larbi, A., Zolezzi, F., Renia, L., Poidinger, M., Naik, S., Newell, E.W., Robson, P., and Ginhoux, F. (2015) Identification of cDC1- and cDC2-committed DC progenitors reveals early lineage priming at the common DC progenitor stage in the bone marrow. Nat. Immunol. 16, 718–728

(14) Grajales-Reyes, G.E., Iwata, A., Albring, J., Wu, X., Tussiwand, R., Kc, W., Kretzer, N.M., Briseño, C.G., Durai, V., Bagadia, P., Haldar, M., Schönheit, J., Rosenbauer, F., Murphy, T.L., and Murphy, K.M. (2015) Batf3 maintains autoactivation of Irf8 for commitment of a CD8α+ conventional DC clonogenic progenitor. Nat. Immunol. 16, 708–717

(15) Bagadia, P., Huang, X., Liu, T.-T., Durai, V., Grajales-Reyes, G.E., Nitschké, M., Modrusan, Z., Granja, J.M., Satpathy, A.T., Briseño, C.G., Gargaro, M., Iwata, A., Kim, S., Chang, H.Y., Shaw, A.S., Murphy, T.L., and Murphy, K.M. (2019) An Nfil3–Zeb2–Id2 pathway imposes Irf8 enhancer switching during cDC1 development. Nat. Immunol. 20, 1174–1185

(16) Minutti, C.M., Piot, C., Pereira da Costa, M., Chakravarty, P., Rogers, N., Huerga Encabo, H., Cardoso, A., Loong, J., Bessou, G., Mionnet, C., Langhorne, J., Bonnet, D., Dalod, M., Tomasello, E., and Reis e Sousa, C. (2024) Distinct ontogenetic lineages dictate cDC2 heterogeneity. Nat. Immunol. 25, 448–461

(17) Rodrigues, P.F., Kouklas, A., Cvijetic, G., Bouladoux, N., Mitrovic, M., Desai, J.V., Lima-Junior, D.S., Lionakis, M.S., Belkaid, Y., Ivanek, R., and Tussiwand, R. (2023) pDC-like cells are pre-DC2 and require KLF4 to control homeostatic CD4 T cells. Sci Immunol. 8, eadd4132

(18) Rodrigues, P.F., Trsan, T., Cvijetic, G., Khantakova, D., Panda, S.K., Liu, Z., Ginhoux, F., Cella, M., and Colonna, M. (2024) Progenitors of distinct lineages shape the diversity of mature type 2 conventional dendritic cells. Immunity. 57, 1567–1585.e1565

(19) Sulczewski, F.B., Maqueda-Alfaro, R.A., Alcántara-Hernández, M., Perez, O.A., Saravanan, S., Yun, T.J., Seong, D., Arroyo Hornero, R., Raquer-McKay, H.M., Esteva, E., Lanzar, Z.R., Leylek, R.A., Adams, N.M., Das, A., Rahman, A.H., Gottfried-Blackmore, A., Reizis, B., and Idoyaga, J. (2023) Transitional dendritic cells are distinct from conventional DC2 precursors and mediate proinflammatory antiviral responses. Nat. Immunol. 24, 1265–1280

(20) Leylek, R., Alcántara-Hernández, M., Lanzar, Z., Lüdtke, A., Perez, O.A., Reizis, B., and Idoyaga, J. (2019) Integrated Cross-Species Analysis Identifies a Conserved Transitional Dendritic Cell Population. Cell Rep. 29, 3736–3750.e3738

(21) Dursun, E., Endele, M., Musumeci, A., Failmezger, H., Wang, S.-H., Tresch, A., Schroeder, T., and Krug, A.B. (2016) Continuous single cell imaging reveals sequential steps of plasmacytoid dendritic cell development from common dendritic cell progenitors. Sci. Rep. 6, 37462

(22) Feng, J., Pucella, J.N., Jang, G., Alcántara-Hernández, M., Upadhaya, S., Adams, N.M., Khodadadi-Jamayran, A., Lau, C.M., Stoeckius, M., Hao, S., Smibert, P., Tsirigos, A., Idoyaga, J., and Reizis, B. (2022) Clonal lineage tracing reveals shared origin of conventional and plasmacytoid dendritic cells. Immunity. 55, 405–422.e411

(23) Onai, N., Kurabayashi, K., Hosoi-Amaike, M., Toyama-Sorimachi, N., Matsushima, K., Inaba, K., and Ohteki, T. (2013) A clonogenic progenitor with prominent plasmacytoid dendritic cell developmental potential. Immunity. 38, 943–957

(24) Dress, R.J., Dutertre, C.-A., Giladi, A., Schlitzer, A., Low, I., Shadan, N.B., Tay, A., Lum, J., Kairi, M.F.B.M., Hwang, Y.Y., Becht, E., Cheng, Y., Chevrier, M., Larbi, A., Newell, E.W., Amit, I., Chen, J., and Ginhoux, F. (2019) Plasmacytoid dendritic cells develop from Ly6D+ lymphoid progenitors distinct from the myeloid lineage. Nat. Immunol. 20, 852–864

(25) Herman, J.S., Sagar, and Grün, D. (2018) FateID infers cell fate bias in multipotent progenitors from single-cell RNA-seq data. Nat. Methods. 15, 379–386

(26) Rodrigues, P.F., Alberti-Servera, L., Eremin, A., Grajales-Reyes, G.E., Ivanek, R., and Tussiwand, R. (2018) Distinct progenitor lineages contribute to the heterogeneity of plasmacytoid dendritic cells. Nat. Immunol. 19, 711–722

(27) Musumeci, A., Lutz, K., Winheim, E., and Krug, A.B. (2019) What Makes a pDC: Recent Advances in Understanding Plasmacytoid DC Development and Heterogeneity. Front. Immunol. 10, 1222

(28) Liu, Z., Wang, H., Li, Z., Dress, R.J., Zhu, Y., Zhang, S., De Feo, D., Kong, W.T., Cai, P., Shin, A., Piot, C., Yu, J., Gu, Y., Zhang, M., Gao, C., Chen, L., Wang, H., Vétillard, M., Guermonprez, P., Kwok, I., Ng, L.G., Chakarov, S., Schlitzer, A., Becher, B., Dutertre, C.A., Su, B., and Ginhoux, F. (2023) Dendritic cell type 3 arises from Ly6C(+) monocyte-dendritic cell progenitors. Immunity. 56, 1761–1777.e1766

(29) Lin, Q., Chauvistré, H., Costa, I.G., Gusmao, E.G., Mitzka, S., Hänzelmann, S., Baying, B., Klisch, T., Moriggl, R., Hennuy, B., Smeets, H., Hoffmann, K., Benes, V., Seré, K., and Zenke, M. (2015) Epigenetic program and transcription factor circuitry of dendritic cell development. Nucl. Acids Res. 43, 9680–9693

(30) Chauvistré, H., and Seré, K. (2020) Epigenetic aspects of DC development and differentiation. Mol. Immunol. 128, 116–124

(31) Ji, Y., Xiao, C., Fan, T., Deng, Z., Wang, D., Cai, W., Li, J., Liao, T., Li, C., and He, J. (2025) The epigenetic hallmarks of immune cells in cancer. Mol. Cancer. 24, 66

(32) Adams, N.M., Galitsyna, A., Tiniakou, I., Esteva, E., Lau, C.M., Reyes, J., Abdennur, N., Shkolikov, A., Yap, G.S., Khodadadi-Jamayran, A., Mirny, L.A., and Reizis, B. (2024) Cohesin-mediated chromatin remodeling controls the differentiation and function of conventional dendritic cells. bioRxiv. 2024.2009.2018.613709

(33) Guak, H., Weiland, M., Ark, A.V., Zhai, L., Lau, K., Corrado, M., Davidson, P., Asiedu, E., Mabvakure, B., Compton, S., DeCamp, L., Scullion, C.A., Jones, R.G., Nowinski, S.M., and Krawczyk, C.M. (2024) Transcriptional programming mediated by the histone demethylase KDM5C regulates dendritic cell population heterogeneity and function. Cell Rep. 43, 114506

(34) De Sá Fernandes, C., Novoszel, P., Gastaldi, T., Krauß, D., Lang, M., Rica, R., Kutschat, A.P., Holcmann, M., Ellmeier, W., Seruggia, D., Strobl, H., and Sibilia, M. (2024) The histone deacetylase HDAC1 controls dendritic cell development and anti-tumor immunity. Cell Rep. 43, 114308

(35) Chauvistré, H., Küstermann, C., Rehage, N., Klisch, T., Mitzka, S., Felker, P., Rose-John, S., Zenke, M., and Seré, K.M. (2014) Dendritic cell development requires histone deacetylase activity. Eur. J. Immunol. 44, 2478–2488

(36) Zhang, Y., Wu, T., He, Z., Lai, W., Shen, X., Lv, J., Wang, Y., and Wu, L. (2023) Regulation of pDC fate determination by histone deacetylase 3. eLife. 12,

(37) Milazzo, G., Mercatelli, D., Di Muzio, G., Triboli, L., De Rosa, P., Perini, G., and Giorgi, F.M. (2020) Histone Deacetylases (HDACs): Evolution, Specificity, Role in Transcriptional Complexes, and Pharmacological Actionability. Genes. 11, 556

(38) Vlaming, H., McLean, C.M., Korthout, T., Alemdehy, M.F., Hendriks, S., Lancini, C., Palit, S., Klarenbeek, S., Kwesi-Maliepaard, E.M., Molenaar, T.M., Hoekman, L., Schmidlin, T.T., Altelaar, A.M., van Welsem, T., Dannenberg, J.H., Jacobs, H., and van Leeuwen, F. (2019) Conserved crosstalk between histone deacetylation and H3K79 methylation generates DOT1L-dose dependency in HDAC1-deficient thymic lymphoma. EMBO J. 38, e101564

(39) Aslam, M.A., Alemdehy, M.F., Kwesi-Maliepaard, E.M., Muhaimin, F.I., Caganova, M., Pardieck, I.N., van den Brand, T., van Welsem, T., de Rink, I., Song, J.-Y., de Wit, E., Arens, R., Jacobs, H., and van Leeuwen, F. (2021) Histone methyltransferase DOT1L controls state-specific identity during B cell differentiation. EMBO Rep. 22, e51184

(40) Scheer, S., Runting, J., Bramhall, M., Russ, B., Zaini, A., Ellemor, J., Rodrigues, G., Ng, J., and Zaph, C. (2020) The Methyltransferase DOT1L Controls Activation and Lineage Integrity in CD4+ T Cells during Infection and Inflammation. Cell Rep. 33, 108505

(41) Kwesi-Maliepaard, E.M., Aslam, M.A., Alemdehy, M.F., van den Brand, T., McLean, C., Vlaming, H., van Welsem, T., Korthout, T., Lancini, C., Hendriks, S., Ahrends, T., van Dinther, D., den Haan, J.M.M., Borst, J., de Wit, E., van Leeuwen, F., and Jacobs, H. (2020) The histone methyltransferase DOT1L prevents antigen-independent differentiation and safeguards epigenetic identity of CD8+ T cells. Proc. Natl. Acad. Sci. U.S.A. 117, 20706–20716

(42) Kealy, L., Di Pietro, A., Hailes, L., Scheer, S., Dalit, L., Groom, J.R., Zaph, C., and Good-Jacobson, K.L. (2020) The Histone Methyltransferase DOT1L Is Essential for Humoral Immune Responses. Cell Rep. 33, 108504

(43) Kealy, L., Runting, J., Thiele, D., and Scheer, S. (2024) An emerging maestro of immune regulation: how DOT1L orchestrates the harmonies of the immune system. Front. Immunol. 15, 1385319

(44) Willemsen, L., Prange, K.H.M., Neele, A.E., van Roomen, C., Gijbels, M., Griffith, G.R., Toom, M.D., Beckers, L., Siebeler, R., Spann, N.J., Chen, H.J., Bosmans, L.A., Gorbatenko, A., van Wouw, S., Zelcer, N., Jacobs, H., van Leeuwen, F., and de Winther, M.P.J. (2022) DOT1L regulates lipid biosynthesis and inflammatory responses in macrophages and promotes atherosclerotic plaque stability. Cell Rep. 41, 111703

(45) Sudholz, H., Schuster, I.S., Foroutan, M., Sng, X., Andoniou, C.E., Doan, A., Camilleri, T., Shen, Z., Zaph, C., Degli-Esposti, M.A., Huntington, N.D., and Scheer, S. (2024) DOT1L maintains NK cell phenotype and function for optimal tumor control. Cell Rep. 43, 114333

(46) Kagoya, Y., Nakatsugawa, M., Saso, K., Guo, T., Anczurowski, M., Wang, C.-H., Butler, M.O., Arrowsmith, C.H., and Hirano, N. (2018) DOT1L inhibition attenuates graft-versus-host disease by allogeneic T cells in adoptive immunotherapy models. Nat. Comm. 9, 1915

(47) Wu, D., Zhang, J., Jun, Y., Liu, L., Huang, C., Wang, W., Yang, C., Xiang, Z., Wu, J., Huang, Y., Meng, D., Yang, Z., Zhou, X., Cheng, C., and Yang, J. (2024) The emerging role of DOT1L in cell proliferation and differentiation: Friend or foe. Histol Histopathol. 39, 425–435

(48) Zhou, Z., Chen, H., Xie, R., Wang, H., Li, S., Xu, Q., Xu, N., Cheng, Q., Qian, Y., Huang, R., Shao, Z., and Xiang, M. (2019) Epigenetically modulated FOXM1 suppresses dendritic cell maturation in pancreatic cancer and colon cancer. Mol. Oncol. 13, 873–893

(49) Tang, Q., Fan, G., Peng, X., Sun, X., Kong, X., Zhang, L., Zhang, C., Liu, Y., Yang, J., Yu, K., Miao, C., Yao, Z., Li, L., Zhang, Z.-S., and Wang, Q. (2025) Gut bacterial L-lysine alters metabolism and histone methylation to drive dendritic cell tolerance. Cell Rep. 44, 115125

(50) Wood, K., Tellier, M., and Murphy, S. (2018) DOT1L and H3K79 Methylation in Transcription and Genomic Stability. Biomolecules. 8,

(51) Vlaming, H., and van Leeuwen, F. (2016) The upstreams and downstreams of H3K79 methylation by DOT1L. Chromosoma. 125, 593–605

(52) Farooq, Z., Banday, S., Pandita, T.K., and Altaf, M. (2016) The many faces of histone H3K79 methylation. Mutat. Res. Rev. Mutat. Res. 768, 46–52

(53) Huff, J.T., Plocik, A.M., Guthrie, C., and Yamamoto, K.R. (2010) Reciprocal intronic and exonic histone modification regions in humans. Nat. Struct. Mol. Biol. 17, 1495–1499

(54) Valencia-Sanchez, M.I., De Ioannes, P., Wang, M., Truong, D.M., Lee, R., Armache, J.P., Boeke, J.D., and Armache, K.J. (2021) Regulation of the Dot1 histone H3K79 methyltransferase by histone H4K16 acetylation. Science. 371,

(55) Ljungman, M., Parks, L., Hulbatte, R., and Bedi, K. (2019) The role of H3K79 methylation in transcription and the DNA damage response. Mutat. Res. Rev. Mutat. Res. 780, 48–54

(56) Wille, C.K., and Sridharan, R. (2022) Connecting the DOTs on Cell Identity. Front. Cell Dev. Biol. 10, 906713

(57) Yu, W., Chory, E.J., Wernimont, A.K., Tempel, W., Scopton, A., Federation, A., Marineau, J.J., Qi, J., Barsyte-Lovejoy, D., Yi, J., Marcellus, R., Iacob, R.E., Engen, J.R., Griffin, C., Aman, A., Wienholds, E., Li, F., Pineda, J., Estiu, G., Shatseva, T., Hajian, T., Al-awar, R., Dick, J.E., Vedadi, M., Brown, P.J., Arrowsmith, C.H., Bradner, J.E., and Schapira, M. (2012) Catalytic site remodelling of the DOT1L methyltransferase by selective inhibitors. Nat. Comm. 3, 1288

(58) Yi, Y., and Ge, S. (2022) Targeting the histone H3 lysine 79 methyltransferase DOT1L in MLL-rearranged leukemias. J. Hemat. Oncol. 15, 35

(59) Perner, F., Berg, T., Sasca, D., Mersiowsky, S.L., Gadrey, J.Y., Thomas, J., Kühn, M.W.M., and Lübbert, M. (2025) Therapeutic targeting of chromatin alterations in leukemia and solid tumors. Int. J. Cancer.

(60) Arnold, O., Barbosa, K., Deshpande, A.J., and Zhu, N. (2022) The Role of DOT1L in Normal and Malignant Hematopoiesis. Front Cell Dev Biol. 10, 917125

(61) Nguyen, V.T.M., Namba, H., Porter, H., Shlyueva, D., Lopez, E., Melcher, A., Beguelin, W., Melnick, A.M., and Helin, K. (2025) Synergistic antitumor effect of combined EZH2 and DOT1L inhibition in B-cell lymphoma. Blood. 145, 2873–2886

(62) Göbel, C., Niccolai, R., de Groot, M.H.P., Jayachandran, J., Traets, J., Kloosterman, D.J., Gregoricchio, S., Morris, B., Kreft, M., Song, J.-Y., Azarang, L., Kasa, E., Oskam, N., de Groot, D., Hoekman, L., Bleijerveld, O.B., Kersten, M.J., Aslam, M.A., van Leeuwen, F., and Jacobs, H. (2025) Targeting DOT1L and EZH2 synergizes in breaking the germinal center identity of diffuse large B-cell lymphoma. Blood. 145, 1802–1813

(63) Borosha, S., Ratri, A., Ghosh, S., Malcom, C.A., Chakravarthi, V.P., Vivian, J.L., Fields, T.A., Rumi, M.A.K., and Fields, P.E. (2022) DOT1L Mediated Gene Repression in Extensively Self-Renewing Erythroblasts. Front. Genet. 13, 828086

(64) Wu, A., Zhi, J., Tian, T., Cihan, A., Cevher, M.A., Liu, Z., David, Y., Muir, T.W., Roeder, R.G., and Yu, M. (2021) DOT1L complex regulates transcriptional initiation in human erythroleukemic cells. Proc. Natl. Acad. Sci. U.S.A. 118,

(65) Malcom, C.A., Ratri, A., Piasecka-Srader, J., Borosha, S., Chakravarthi, V.P., Alvarez, N.S., Vivian, J.L., Fields, T.A., Karim Rumi, M.A., and Fields, P.E. (2021) Primitive Erythropoiesis in the Mouse is Independent of DOT1L Methyltransferase Activity. Front. Cell Dev. Biol. 9, 813503

(66) Cao, K., Ugarenko, M., Ozark, P.A., Wang, J., Marshall, S.A., Rendleman, E.J., Liang, K., Wang, L., Zou, L., Smith, E.R., Yue, F., and Shilatifard, A. (2020) DOT1L-controlled cell-fate determination and transcription elongation are independent of H3K79 methylation. Proc. Natl. Acad. Sci. U.S.A. 117, 27365–27373

(67) van Welsem, T., Korthout, T., Ekkebus, R., Morais, D., Molenaar, T.M., van Harten, K., Poramba-Liyanage, D.W., Sun, S.M., Lenstra, T.L., Srivas, R., Ideker, T., Holstege, F.C.P., van Attikum, H., El Oualid, F., Ovaa, H., Stulemeijer, I.J.E., Vlaming, H., and van Leeuwen, F. (2018) Dot1 promotes H2B ubiquitination by a methyltransferase-independent mechanism. Nucl. Acids Res. 46, 11251–11261

(68) Stulemeijer, I.J.E., Pike, B.L., Faber, A.W., Verzijlbergen, K.F., van Welsem, T., Frederiks, F., Lenstra, T.L., Holstege, F.C., Gasser, S.M., and van Leeuwen, F. (2011) Dot1 binding induces chromatin rearrangements by histone methylation-dependent and -independent mechanisms. Epigen. Chrom. 4, 2

(69) Hameyer, D., Loonstra, A., Eshkind, L., Schmitt, S., Antunes, C., Groen, A., Bindels, E., Jonkers, J., Krimpenfort, P., Meuwissen, R., Rijswijk, L., Bex, A., Berns, A., and Bockamp, E. (2007) Toxicity of ligand-dependent Cre recombinases and generation of a conditional Cre deleter mouse allowing mosaic recombination in peripheral tissues. Physiol. Genomics. 31, 32–41

(70) Skarnes, W.C., Rosen, B., West, A.P., Koutsourakis, M., Bushell, W., Iyer, V., Mujica, A.O., Thomas, M., Harrow, J., Cox, T., Jackson, D., Severin, J., Biggs, P., Fu, J., Nefedov, M., de Jong, P.J., Stewart, A.F., and Bradley, A. (2011) A conditional knockout resource for the genome-wide study of mouse gene function. Nature. 474, 337–342

(71) Ashburner, M., Ball, C.A., Blake, J.A., Botstein, D., Butler, H., Cherry, J.M., Davis, A.P., Dolinski, K., Dwight, S.S., Eppig, J.T., Harris, M.A., Hill, D.P., Issel-Tarver, L., Kasarskis, A., Lewis, S., Matese, J.C., Richardson, J.E., Ringwald, M., Rubin, G.M., and Sherlock, G. (2000) Gene Ontology: tool for the unification of biology. Nat. Genet. 25, 25–29

(72) The Gene Ontology Consortium, and Aleksander, S.A., and Balhoff, J., and Carbon, S., and Cherry, J.M., and Drabkin, H.J., and Ebert, D., and Feuermann, M., and Gaudet, P., and Harris, N.L., and Hill, D.P., and Lee, R., and Mi, H., and Moxon, S., and Mungall, C.J., and Muruganugan, A., and Mushayahama, T., and Sternberg, P.W., and Thomas, P.D., and Van Auken, K., and Ramsey, J., and Siegele, D.A., and Chisholm, R.L., and Fey, P., and Aspromonte, M.C., and Nugnes, M.V., and Quaglia, F., and Tosatto, S., and Giglio, M., and Nadendla, S., and Antonazzo, G., and Attrill, H., and dos Santos, G., and Marygold, S., and Strelets, V., and Tabone, C.J., and Thurmond, J., and Zhou, P., and Ahmed, S.H., and Asanitthong, P., and Luna Buitrago, D., and Erdol, M.N., and Gage, M.C., and Ali Kadhum, M., and Li, K.Y.C., and Long, M., and Michalak, A., and Pesala, A., and Pritazahra, A., and Saverimuttu, S.C.C., and Su, R., and Thurlow, K.E., and Lovering, R.C., and Logie, C., and Oliferenko, S., and Blake, J., and Christie, K., and Corbani, L., and Dolan, M.E., and Drabkin, H.J., and Hill, D.P., and Ni, L., and Sitnikov, D., and Smith, C., and Cuzick, A., and Seager, J., and Cooper, L., and Elser, J., and Jaiswal, P., and Gupta, P., and Jaiswal, P., and Naithani, S., and Lera-Ramirez, M., and Rutherford, K., and Wood, V., and De Pons, J.L., and Dwinell, M.R., and Hayman, G.T., and Kaldunski, M.L., and Kwitek, A.E., and Laulederkind, S.J.F., and Tutaj, M.A., and Vedi, M., and Wang, S.-J., and D’Eustachio, P., and Aimo, L., and Axelsen, K., and Bridge, A., and Hyka-Nouspikel, N., and Morgat, A., and Aleksander, S.A., and Cherry, J.M., and Engel, S.R., and Karra, K., and Miyasato, S.R., and Nash, R.S., and Skrzypek, M.S., and Weng, S., and Wong, E.D., and Bakker, E., and Berardini, T.Z., and Reiser, L., and Auchincloss, A., and Axelsen, K., and Argoud-Puy, G., and Blatter, M.-C., and Boutet, E., and Breuza, L., and Bridge, A., and Casals-Casas, C., and Coudert, E., and Estreicher, A., and Livia Famiglietti, M., and Feuermann, M., and Gos, A., and Gruaz-Gumowski, N., and Hulo, C., and Hyka-Nouspikel, N., and Jungo, F., and Le Mercier, P., and Lieberherr, D., and Masson, P., and Morgat, A., and Pedruzzi, I., and Pourcel, L., and Poux, S., and Rivoire, C., and Sundaram, S., and Bateman, A., and Bowler-Barnett, E., and Bye-A-Jee, H., and Denny, P., and Ignatchenko, A., and Ishtiaq, R., and Lock, A., and Lussi, Y., and Magrane, M., and Martin, M.J., and Orchard, S., and Raposo, P., and Speretta, E., and Tyagi, N., and Warner, K., and Zaru, R., and Diehl, A.D., and Lee, R., and Chan, J., and Diamantakis, S., and Raciti, D., and Zarowiecki, M., and Fisher, M., and James-Zorn, C., and Ponferrada, V., and Zorn, A., and Ramachandran, S., and Ruzicka, L., and Westerfield, M. (2023) The Gene Ontology knowledgebase in 2023. Genetics. 224,

(73) Ge, S.X., Jung, D., and Yao, R. (2019) ShinyGO: a graphical gene-set enrichment tool for animals and plants. Bioinformatics. 36, 2628–2629

(74) Malik, M., de Leeuw, W.-J., Aslam, M.A., Kwesi-Maliepaard, E.M., van den Brand, T., van den Broek, B., Kempers, M., Hoekman, L., Proost, N., van Welsem, T., de Wit, E., Borst, J., Jacobs, H., and van Leeuwen, F. (2025) Histone methyltransferase DOT1L maintains identity and restricts cytotoxic potential of CD8 T cells. bioRxiv. 2025.2001.2020.633937

(75) Szklarczyk, D., Kirsch, R., Koutrouli, M., Nastou, K., Mehryary, F., Hachilif, R., Gable, A.L., Fang, T., Doncheva, Nadezhda T., Pyysalo, S., Bork, P., Jensen, Lars J., and von Mering, C. (2022) The STRING database in 2023: protein–protein association networks and functional enrichment analyses for any sequenced genome of interest. Nucl. Acids Res. 51, D638–D646

(76) Steger, D.J., Lefterova, M.I., Ying, L., Stonestrom, A.J., Schupp, M., Zhuo, D., Vakoc, A.L., Kim, J.E., Chen, J., Lazar, M.A., Blobel, G.A., and Vakoc, C.R. (2008) DOT1L/KMT4 recruitment and H3K79 methylation are ubiquitously coupled with gene transcription in mammalian cells. Mol. Cell. Biol. 28, 2825–2839

(77) Lutz, M.B., Ali, S., Audiger, C., Autenrieth, S.E., Berod, L., Bigley, V., Cyran, L., Dalod, M., Dörrie, J., Dudziak, D., Flórez-Grau, G., Giusiano, L., Godoy, G.J., Heuer, M., Krug, A.B., Lehmann, C.H.K., Mayer, C.T., Naik, S.H., Scheu, S., Schreibelt, G., Segura, E., Seré, K., Sparwasser, T., Tel, J., Xu, H., and Zenke, M. (2023) Guidelines for mouse and human DC generation. Eur. J. Immunol. 53, e2249816

(78) Clausen, B.E., Amon, L., Backer, R.A., Berod, L., Bopp, T., Brand, A., Burgdorf, S., Chen, L., Da, M., Distler, U., Dress, R.J., Dudziak, D., Dutertre, C.A., Eich, C., Gabele, A., Geiger, M., Ginhoux, F., Giusiano, L., Godoy, G.J., Hamouda, A.E.I., Hatscher, L., Heger, L., Heidkamp, G.F., Hernandez, L.C., Jacobi, L., Kaszubowski, T., Kong, W.T., Lehmann, C.H.K., Lopez-Lopez, T., Mahnke, K., Nitsche, D., Renkawitz, J., Reza, R.A., Saez, P.J., Schlautmann, L., Schmitt, M.T., Seichter, A., Sielaff, M., Sparwasser, T., Stoitzner, P., Tchitashvili, G., Tenzer, S., Tochoedo, N.R., Vurnek, D., Zink, F., and Hieronymus, T. (2023) Guidelines for mouse and human DC functional assays. Eur. J. Immunol. 53, e2249925

(79) Chory, E.J., Calarco, J.P., Hathaway, N.A., Bell, O., Neel, D.S., and Crabtree, G.R. (2019) Nucleosome Turnover Regulates Histone Methylation Patterns over the Genome. Mol. Cell. 73, 61–72.e63

(80) Liu, C., Yang, Q., Zhu, Q., Lu, X., Li, M., Hou, T., Li, Z., Tang, M., Li, Y., Wang, H., Yang, Y., Wang, H., Zhao, Y., Wen, H., Liu, X., Mao, Z., and Zhu, W.G. (2020) CBP mediated DOT1L acetylation confers DOT1L stability and promotes cancer metastasis. Theranostics. 10, 1758–1776

(81) Song, T., Zou, Q., Yan, Y., Lv, S., Li, N., Zhao, X., Ma, X., Liu, H., Tang, B., and Sun, L. (2021) DOT1L O-GlcNAcylation promotes its protein stability and MLL-fusion leukemia cell proliferation. Cell Rep. 36, 109739

(82) De Vos, D., Frederiks, F., Terweij, M., van Welsem, T., Verzijlbergen, K.F., Iachina, E., de Graaf, E.L., Altelaar, A.F.M., Oudgenoeg, G., Heck, A.J., Krijgsveld, J., Bakker, B.M., and van Leeuwen, F. (2011) Progressive methylation of ageing histones by Dot1 functions as a timer. EMBO Rep. 12, 956–962

(83) Rodrigues, P.F., and Tussiwand, R. (2020) Novel concepts in plasmacytoid dendritic cell (pDC) development and differentiation. Mol. Immunol. 126, 25–30

(84) Valente, M., Collinet, N., Vu Manh, T.P., Popoff, D., Rahmani, K., Naciri, K., Bessou, G., Rua, R., Gil, L., Mionnet, C., Milpied, P., Tomasello, E., and Dalod, M. (2023) Novel mouse models based on intersectional genetics to identify and characterize plasmacytoid dendritic cells. Nat Immunol. 24, 714–728

(85) Liu, T.T., Kim, S., Desai, P., Kim, D.H., Huang, X., Ferris, S.T., Wu, R., Ou, F., Egawa, T., Van Dyken, S.J., Diamond, M.S., Johnson, P.F., Kubo, M., Murphy, T.L., and Murphy, K.M. (2022) Ablation of cDC2 development by triple mutations within the Zeb2 enhancer. Nature. 607, 142–148

(86) Ding, L., and Morrison, S.J. (2013) Haematopoietic stem cells and early lymphoid progenitors occupy distinct bone marrow niches. Nature. 495, 231–235

(87) Mooney, C.J., Cunningham, A., Tsapogas, P., Toellner, K.-M., and Brown, G. (2017) Selective Expression of Flt3 within the Mouse Hematopoietic Stem Cell Compartment. Int. J. Mol. Sci. 18,

(88) Liu, K., Waskow, C., Liu, X., Yao, K., Hoh, J., and Nussenzweig, M. (2007) Origin of dendritic cells in peripheral lymphoid organs of mice. Nat. Immunol. 8, 578–583

(89) Zhu, Y., Cai, P., Li, Z., Zhang, S., Kong, W.T., Wang, H., Gao, F., Zeng, Y., Qian, J., Su, B., Liu, Z., and Ginhoux, F. (2025) Transcription factors TCF4 and KLF4 respectively control the development of the DC2A and DC2B lineages. Nat. Immunol.,

(90) Kapahnke, K., Plenge, T., Klaus, T., Gupta, M.K., Anand, D., Onder, T.T., Perner, B., Schnoder, T.M., Thol, F.R., Damm, F., Heidel, F.H., and Perner, F. (2025) The histone-methyltransferase DOT1L cooperates with LSD1 to control cell division in blast-phase MPN. Leukemia.

(91) Kwesi-Maliepaard, E.M., Malik, M., van Welsem, T., van Doorn, R., Vermeer, M.H., Vlaming, H., Jacobs, H., and van Leeuwen, F. (2022) DOT1L inhibition does not modify the sensitivity of cutaneous T cell lymphoma to pan-HDAC inhibitors in vitro. Front. Genet. 13, 1032958

(92) Kasuga, Y., Ouda, R., Watanabe, M., Sun, X., Kimura, M., Hatakeyama, S., and Kobayashi, K.S. (2023) FBXO11 constitutes a major negative regulator of MHC class II through ubiquitin-dependent proteasomal degradation of CIITA. Proc. Natl. Acad. Sci. U.S.A. 120, e2218955120

(93) Radman-Livaja, M., Verzijlbergen, K.F., Weiner, A., van Welsem, T., Friedman, N., Rando, O.J., and van Leeuwen, F. (2011) Patterns and mechanisms of ancestral histone protein inheritance in budding yeast. PLoS Biol. 9, e1001075

(94) Katan-Khaykovich, Y., and Struhl, K. (2005) Heterochromatin formation involves changes in histone modifications over multiple cell generations. EMBO J. 24, 2138–2149-2149

(95) Zee, B.M., Levin, R.S., Xu, B., LeRoy, G., Wingreen, N.S., and Garcia, B.A. (2010) In vivo residue-specific histone methylation dynamics. J. Biol. Chem. 285, 3341–3350

(96) Sweet, S.M.M., Li, M., Thomas, P.M., Durbin, K.R., and Kelleher, N.L. (2010) Kinetics of re-establishing H3K79 methylation marks in global human chromatin. J. Biol. Chem. 285, 32778–32786

(97) Scharf, A.N.D., Barth, T.K., and Imhof, A. (2009) Establishment of histone modifications after chromatin assembly. Nucl. Acids Res. 37, 5032–5040

(98) Alabert, C., Barth, T.K., Reverón-Gómez, N., Sidoli, S., Schmidt, A., Jensen, O.N., Imhof, A., and Groth, A. (2015) Two distinct modes for propagation of histone PTMs across the cell cycle. Genes Dev. 29, 585–590

(99) Patel, A.A., Ginhoux, F., and Yona, S. (2021) Monocytes, macrophages, dendritic cells and neutrophils: an update on lifespan kinetics in health and disease. Immunology. 163, 250–261

(100) van den Elsen, P.J. (2011) Expression regulation of major histocompatibility complex class I and class II encoding genes. Front. Immunol. 2, 48

(101) Kanojia, A., Roy, G., Madhubala, R., and Muthuswami, R. (2024) Interplay between DOT1L and HDAC1 regulates Leishmania donovani infection in human THP-1 cells. Acta Trop. 258, 107352

