## Supplementary Figures for "Histone methyltransferase DOT1L differentially affects the development of dendritic cell subsets"

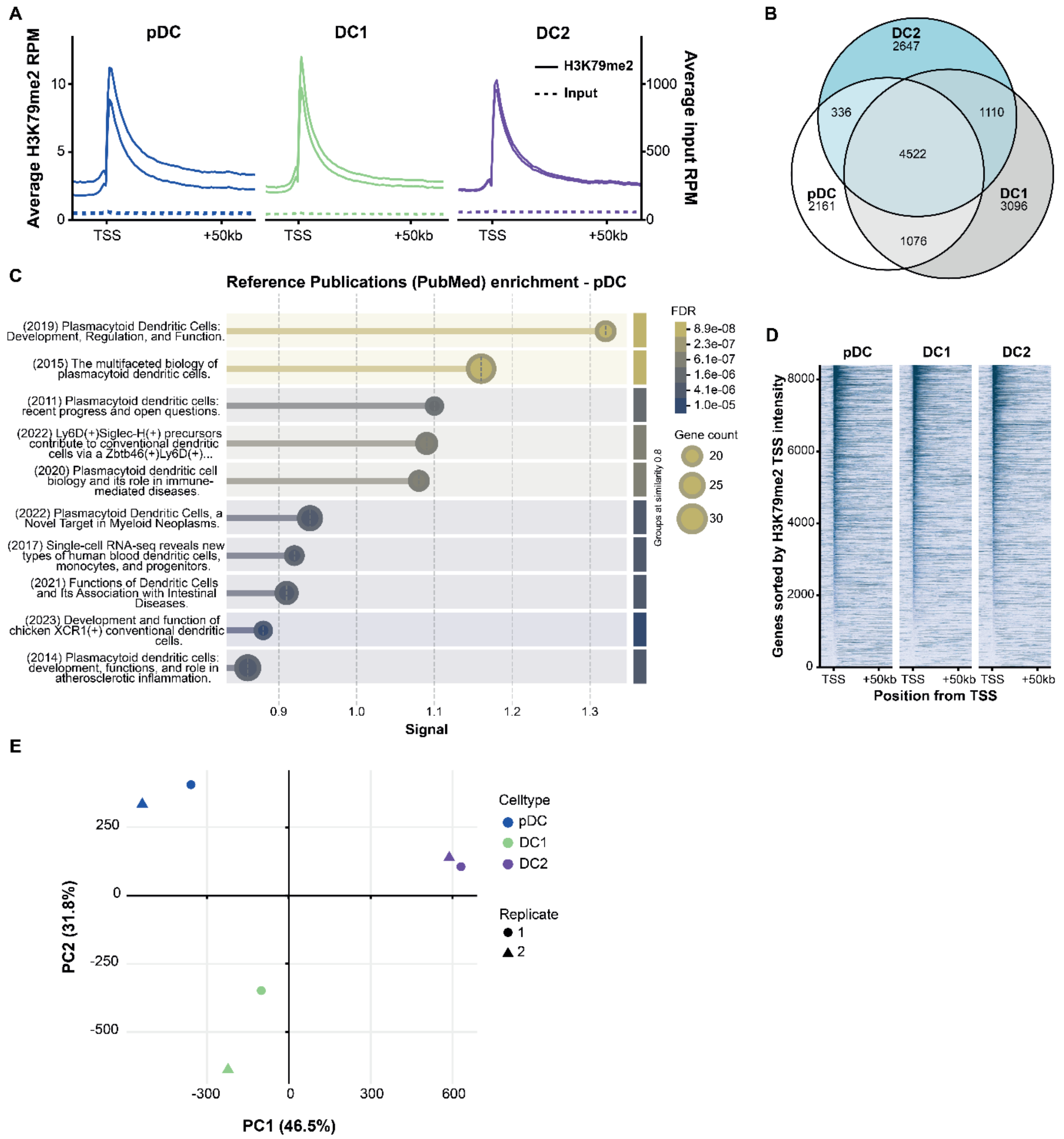


### Supplementary Figure S1 – ChIPseq reveals distinct H3K79 methylation patterns between DC subsets

Siglec-H$^{pos}$ BST2$^{pos}$ pDCs, CD11c$^{pos}$ XCR1$^{pos}$ cDC1s and CD11c$^{pos}$ sirpα$^{pos}$ cDC2s were sorted from the bone-marrow (BM) of *Dot1l-WT* mice, immunoprecipitated for H3K79me2, peaks were identified by sequencing. **(A)** Meta-gene plots quantifying the H3K79me2 signal near the transcriptional start site (TSS) in the sorted populations. **(B)** Venn diagram showing the number of quantified peaks per cell-type and their overlap (aggregate of N=2 biological replicates). **(C)** Gene set enrichment analysis of peaks uniquely mapped to pDCs, searched against the PubMed database using String {Szklarczyk, 2022 #85}. Biological replicates are plotted individually. **(D)** Tornadoplot visualizing H3K79me2 signal near the TSS (up to +2kb) of the top 50% genes based on expression (RNAseq). Genes are ranked from high signal to low, data shown are aggregates of N=2 biological replicates per sorted DC subset. **(E)** Principal component analysis (PCA) of **(D)** highlighting both biological replicates from each subset.

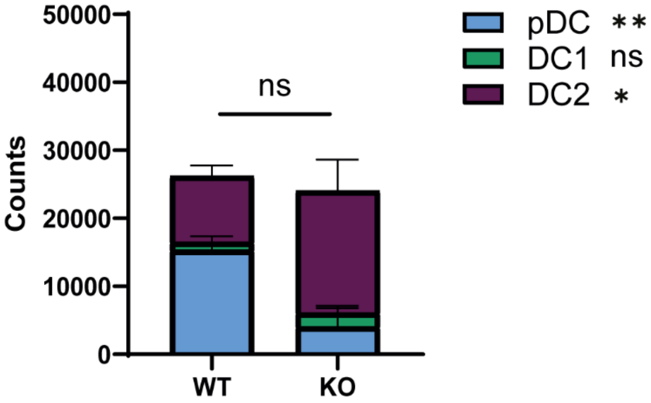


**Supplementary Figure S2 – *In vitro* deletion of *Dot1l* in FLT3L- and SCF- supplemented BM cultures did not affect the number of viable cells**

Total BM of Cre-$ER^{T2}$ $Dot1l^{wt/wt}$ and Cre-$ER^{T2}$ $Dot1l^{fl/fl}$ mice was cultured with or without 4-hydroxytamoxifen and with SCF and FLT3L to facilitate the development of DCs. On day 7, viable cell counts were determined. pDCs were gated as $BST2^{pos}$ Siglec-$H^{pos}$, cDC2s as $CD11c^{pos}$ MHC class $II^{pos}$ $Sirp\alpha^{pos}$ and cDC1s as $CD11c^{pos}$ MHC class $II^{pos}$ $XCR1^{pos}$. Depicted are the number of viable cells from each DC subset per 100,000 total viable cells in the culture, comparing *Dot1l* WT (Cre-$ER^{T2}$;*Dot1l*$^{wt/wt}$ + tamoxifen) and *Dot1l* KO (Cre-$ER^{T2}$;*Dot1l*$^{fl/fl}$ + tamoxifen). Plotted in the graph, statistics represent the difference in total number of cells between WT versus KO conditions. Shown next to the legend, statistics represent the difference in total number of cells between WT versus KO conditions per subset. A total of three independent experiments was performed. Error bars indicate mean ± SD. *p < 0.05**p < 0.01, ***p < 0.001, ****p < 0.0001.

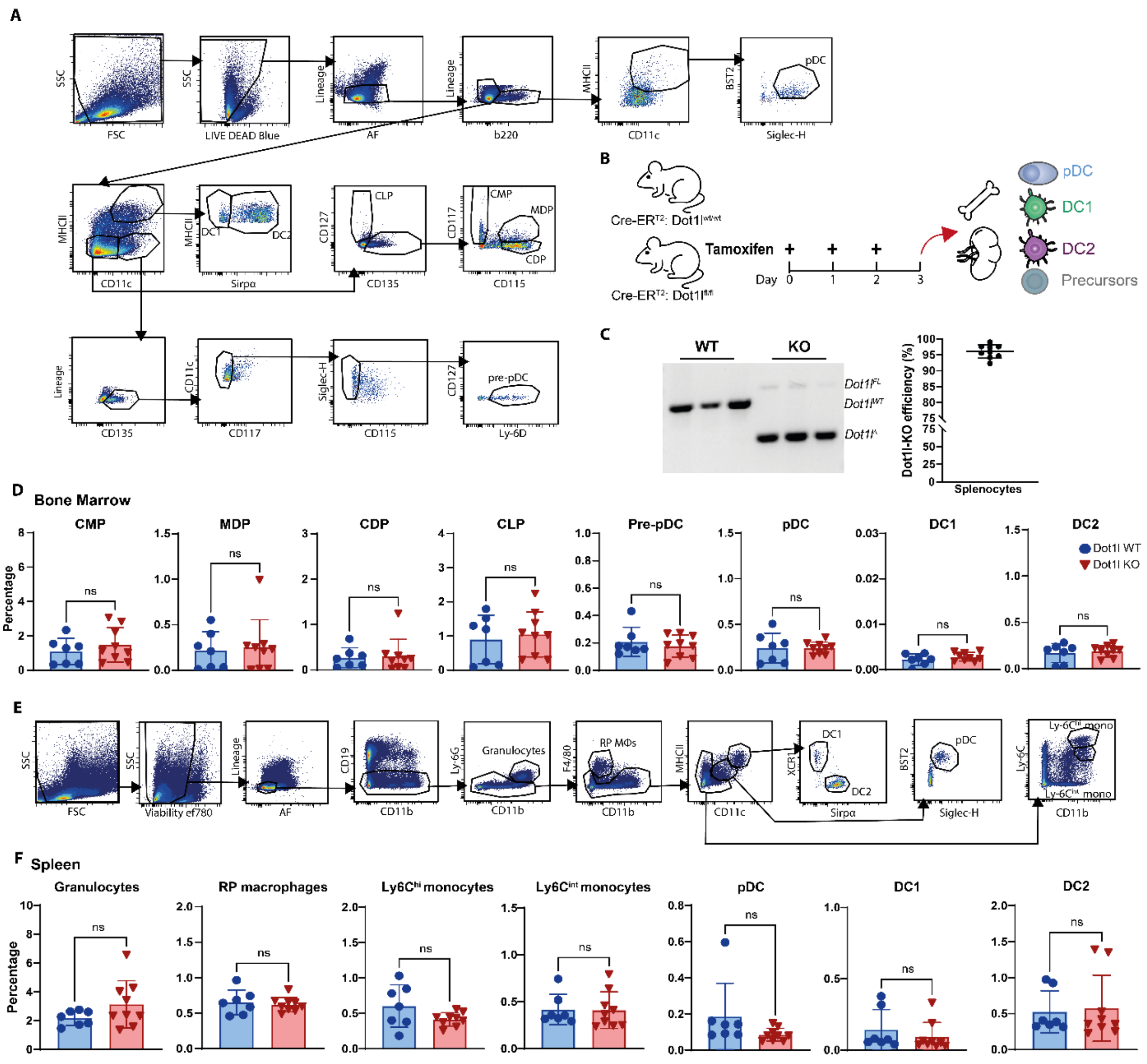


### Supplementary Figure S3 – Gating strategy and population percentages in the BM and spleen on day 3 after *in vivo* deletion of *Dot1l*

**(A)** Overview of the gating strategy to identify lymphoid and myeloid precursors, as well as mature DC subsets from BM. **(B)** Cre-ER$^{T2}$*Dot1l*$^{wt/wt}$ and Cre-ER$^{T2}$*Dot1l*$^{fl/fl}$ mice were injected three times with tamoxifen. Subsequently, BM and spleen were harvested, and several precursors and mature cell subsets were evaluated. **(C)** Genomic DNA was isolated and a region surrounding floxed exon 2 of *Dot1L* was amplified by PCR (left; N=3, representative replicates). The frequency of WT (*Dot1l*$^{WT}$), floxed (*Dot1l*$^{FL}$) and floxed out (*Dot1l*[illegible]) alleles was quantified by ImageJ, after correcting for product size (right; N=9). **(D)** Myeloid and lymphoid precursor percentages of Live Lineage$^{neg}$ AF$^{neg}$ cells in *Dot1l* WT (Cre-ER$^{T2}$*Dot1l*$^{wt/wt}$) and *Dot1l* KO (Cre-ER$^{T2}$ *Dot1l*$^{fl/fl}$) BM on day 3 after 3 tamoxifen injections. **(E)** Overview of the gating strategy to identify mature cell subsets in the spleen. **(F)** Splenic cell subset percentages of Live Lineage$^{neg}$ AF$^{neg}$ cells in *Dot1l* WT (Cre-ER$^{T2}$ Dot1l$^{wt/wt}$) and *Dot1l* KO (Cre-ER$^{T2}$ Dot1l$^{fl/fl}$) spleen on day 3 after 3 tamoxifen injections. The individual dots represent individual mice with a total of three independent experiments performed. Error bars indicate mean ± SD. *$p < 0.05$, **$p < 0.01$, ***$p < 0.001$, ****$p < 0.0001$.

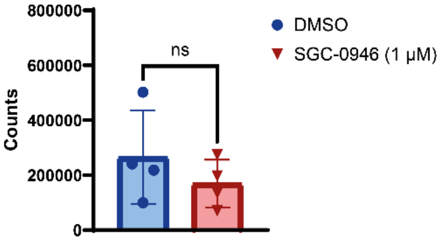


**Supplementary Figure S4 – *In vitro* inhibition of DOT1L in FLT3L- and SCF- supplemented BM cultures does not affect the number of viable cells**

Total BM of WT C57BL/6 mice was cultured with SCF and FLT3L to facilitate the development of DCs. Shown are viable cell counts of DC subsets in cultures without inhibitor (0.1% DMSO as vehicle control) compared to cultures with inhibitor SGC-0946 (1 μM). DOT1L inhibitor SGC-0946 was added at day 0, 3 and 5 to ensure constant inhibition of DOT1L during DC development. Viable cell counts were calculated based on the total amount of cells per sample. The individual dots represent the average of technical replicates in one independent experiment, with a total of three independent experiments performed. Error bars indicate mean ± SD. $^*p < 0.05^{**}p < 0.01$, $^{***}p < 0.001$, $^{****}p < 0.0001$.

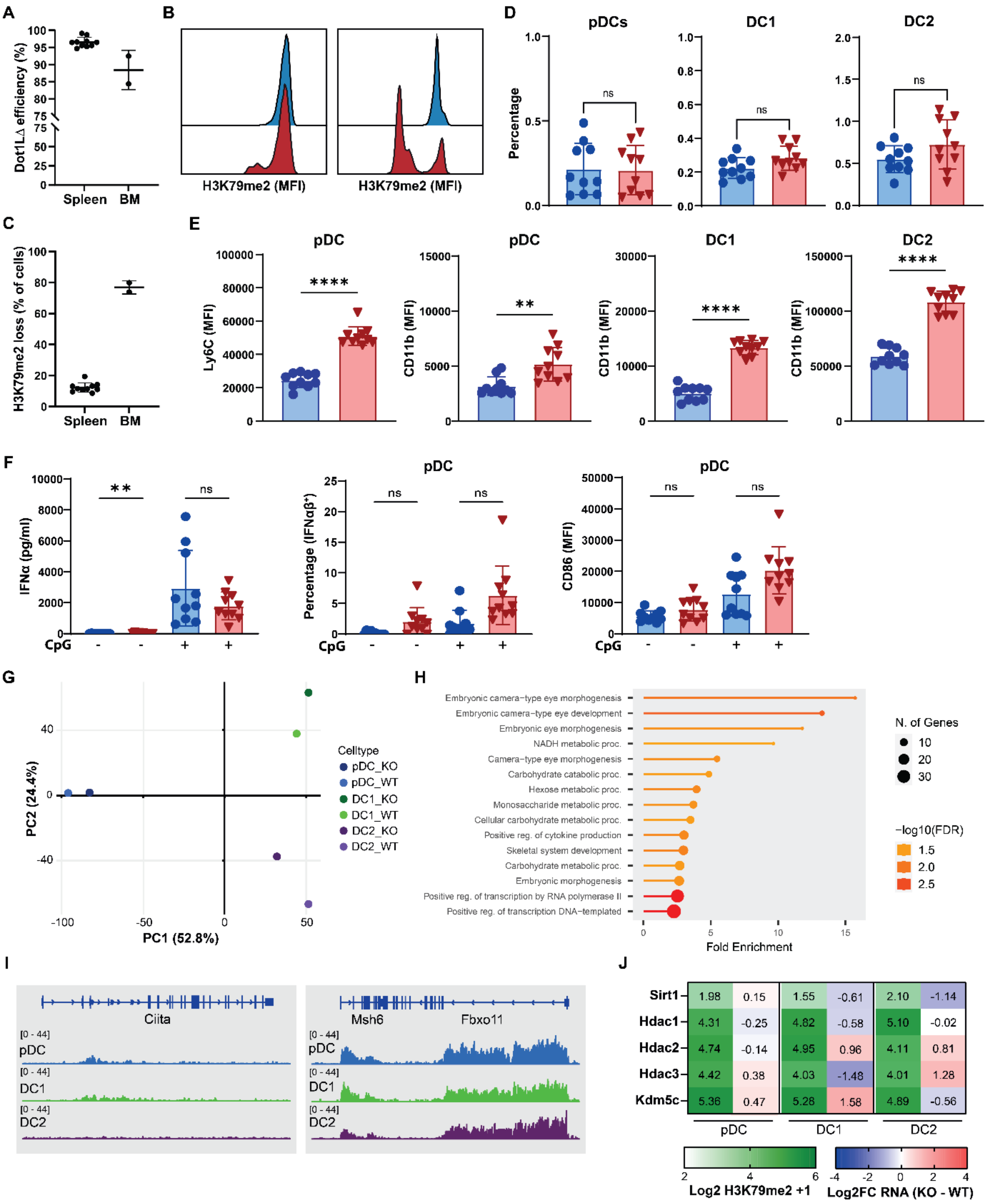


### Supplementary Figure S5 – Characterization of DCs following *in vivo* deletion of *Dot1l* in the spleen and BM

**(A)** Cre-ER$^{T2}$ *Dot1l*$^{wt/wt}$ and Cre-ER$^{T2}$ *Dot1l*$^{fl/fl}$ mice were injected with tamoxifen on three subsequent days to induce KO of *Dot1l*. On day 12, organs were harvested, and several (functional) experiments were performed on the splenocytes and BM cells. The region surrounding exon 2 of *Dot1L* was amplified and the frequency of *Dot1l*$^{Δ}$ alleles was quantified as described previously (BM = bone marrow). **(B)** Histograms of H3K79me2 in representative *Dot1l* WT and KO samples derived from either spleen (left) or BM (right). For the KO samples, cells overlapping with the WT peak were classified as H3K79me2 high, and the remainder as H3K79me2 low. **(C)** Quantification of the H3K79me2 low fractions from S5B (N=10 for spleen, N=2 (pooled replicates) for BM). **(D)** Splenic DC subset percentages of Live Lineage$^{neg}$ AF$^{neg}$ cells in *Dot1l* WT (Cre-ER$^{T2}$ Dot1l$^{wt/wt}$) and *Dot1l* KO (Cre-ER$^{T2}$ Dot1l$^{fl/fl}$) conditions. pDCs were gated as BST2$^{pos}$ Siglec-H$^{pos}$, cDC2s as CD11c$^{pos}$ MHC class II$^{pos}$ Sirpα$^{pos}$ and cDC1s as CD11c$^{pos}$ MHC class II$^{pos}$ XCR1$^{pos}$. **(E)** Quantification of specific markers on pDCs and cDCs that were differentially expressed between *Dot1l* WT and *Dot1l* KO conditions based on the MFI of (N=10) biological replicates. **(F)** On day 12, splenocytes were stimulated with Class A CpGs (ODN1585). IFNα levels were determined in the supernatant using ELISA. Additionally, cells were stained intracellularly for IFNαβ. After overnight stimulation, cells were also stained for maturation markers. Conditions are shown with and without the addition of CpGs. **(G)** Principal component analysis (PCA) of pooled and sorted DC subsets (N=1) from both WT and Dot1l KO mice. **(H)** Pie-chart (left) highlighting the number of down- and up-regulated genes in all KO populations (based on Log2FC <-0.5 and >0.5 respectively); (right) GO-enrichment of genes that were downregulated in expression (Log2 <-0.5) in all sorted subsets, ranked by fold-enrichment and visualized using ShinyGO. **(I)** Representative H3K79me2 ChIP-seq tracks for *Ciita* and *Fbxo11* exported from IGV. Data scales were standardized to visualize quantitative differences both between genes and between DC subsets. **(J)** Heatmap of selected epigenetic modifiers showing H3K79me2 signal near the TSS (N=2, green gradient) and expression differences between Dot1l-KO and WT (N=1, blue-red gradient) in the sorted DC subsets. Error bars indicate mean ± SD. *p < 0.05, **p < 0.01, ***p < 0.001, ****p < 0.0001. Error bars indicate mean ± SD. *p < 0.05, **p < 0.01, ***p < 0.001, ****p < 0.0001.
